# Data-driven synapse classification reveals a logic of glutamate receptor diversity

**DOI:** 10.1101/2024.12.11.628056

**Authors:** Kristina D. Micheva, Anish K. Simhal, Jenna Schardt, Stephen J Smith, Richard J. Weinberg, Scott F. Owen

## Abstract

The rich diversity of synapses facilitates the capacity of neural circuits to transmit, process and store information. We used multiplex super-resolution proteometric imaging through array tomography to define features of single synapses in mouse neocortex. We find that glutamatergic synapses cluster into subclasses that parallel the distinct biochemical and functional categories of receptor subunits: GluA1/4, GluA2/3 and GluN1/GluN2B. Two of these subclasses align with physiological expectations based on synaptic plasticity: large AMPAR-rich synapses may represent potentiated synapses, whereas small NMDAR-rich synapses suggest “silent” synapses. The NMDA receptor content of large synapses correlates with spine neck diameter, and thus the potential for coupling to the parent dendrite. Overall, ultrastructural features predict receptor content of synapses better than parent neuron identity does, suggesting synapse subclasses act as fundamental elements of neuronal circuits. No barriers prevent future generalization of this approach to other species, or to study of human disorders and therapeutics.

## Introduction

Synapses display remarkable diversity in size, molecular composition, subcellular localization, and function^1–13^ thus shaping the transmission and processing of information in neural circuits. Active regulation of synaptic strength associated with modifications in the size and composition of receptors is fundamental to memory storage and neuronal plasticity^14–29^.

Even within a single tightly-restricted anatomical region, the size of individual synaptic contacts varies by at least a factor of ten, as does unitary transmission strength^1,2,4–6,8^. The molecular composition of synapses is tightly orchestrated throughout brain development^30–36^, and risk genes for prevalent neurological disorders including autism, epilepsy, and psychosis are heavily weighted towards synaptic function^37–42^. Quantitative, single-synapse-level profiling to define the logic underlying these synapse populations is therefore critical to understanding both normal and disordered brain development and function.

Here we introduce an approach for single-synapse population profiling based on conjugate array tomography (AT), which uses immunofluorescence followed by scanning EM on serial ultrathin sections to reveal both protein content and tissue ultrastructure^43^. We apply conjugate AT to adult mouse temporal association cortex (TeA). Array tomography has unmatched capacity to bring both multiplexed super-resolution volume fluorescence and volume electron microscopy to bear on individual synapses^10,43–51^, but has not yet been fully exploited for high- dimensional microproteometric analysis. Multi-channel immunofluorescence images enable quantitative sampling of the protein composition of single synapses, while electron microscopy allows precise quantification of the size, shape, and ultrastructural context of the same individual synapses. Machine-learning tools efficiently extract mechanistic insight from this rich multi- dimensional data.

Of the hundreds of distinct proteins present at individual synapses^37,52–55^, the proteins that most directly impact synaptic strength include neurotransmitter receptors and postsynaptic density scaffolds^12,53,56–59^. We therefore selected a set of proteins based on their direct role in generation of excitatory postsynaptic currents, and their relevance to activity-dependent plasticity and thus to memory storage^14,15,23,54^. These include four AMPA-type and two NMDA-type glutamate receptor subunits, along with the scaffold proteins PSD-95, gephyrin and synapsin 1/2, and the cell-type markers GABA and vesicular glutamate transporter 1. The tight three-dimensional registration of volume immunofluorescence and electron microscopy images in our data reveals important connections between ultrastructural and molecular features of individual synapses, and sheds new light on rules governing the scaling of glutamate receptor content with synapse size and target cell identity.

Our analysis indicates that the myriad of diverse glutamatergic synapses in our neocortical target region can be clustered into a modest number of subclasses based on glutamate receptor expression. This clustering is robust even when only multi-channel immunofluorescence data is available, suggesting that this synapse classification rubric can be applied across experimental systems and areas of investigation. The logic of these subclasses can be readily interpreted in the context of distinct physiological roles. For example, large AMPAR-rich synapses suggest full potentiation, while small NMDAR-rich synapses that lack detectable AMPA receptors suggest functionally “silent” synapses. Volume electron microscopy reveals that ultrastructural features like spine head and neck diameter are better predictors of the proteometric content of individual synapses than pre- or postsynaptic parent neuron identity.

This method to profile the diversity of synapses promises to advance our basic understanding of brain development, circuit function, learning and memory. Furthermore, by providing a quantitative rubric to characterize deficits in specific populations of synapses, we offer a valuable new tool to elucidate the synaptic basis of brain and memory disorders.

## Results

### Conjugate array tomography enables multiplex immunolabeling of ultrastructurally- identified synapses

We used multiplex immunolabeling and conjugate AT to explore the receptor composition and morphology of glutamatergic synapses in layers 2/3 of temporal association cortex (TeA) of an adult mouse (Fig.1a-c). Here we present a conjugate “IF-SEM” dataset with co-registered immunofluorescence and electron microscopy, and a larger immunofluorescence-only “IF-only” dataset that does not include electron microscopy. The conjugate “IF-SEM” dataset is a subvolume of the “IF-only” dataset. We used 4 cycles of immunolabeling to localize synapses and quantify protein content (Table 1), choosing synapsin 1/2 as a general presynaptic marker and PSD-95 as a postsynaptic marker for glutamatergic synapses. Inhibitory synapses were distinguished by colocalization of GABA (presynaptic) and gephyrin (postsynaptic) (Fig.1d).

**Figure 1.**
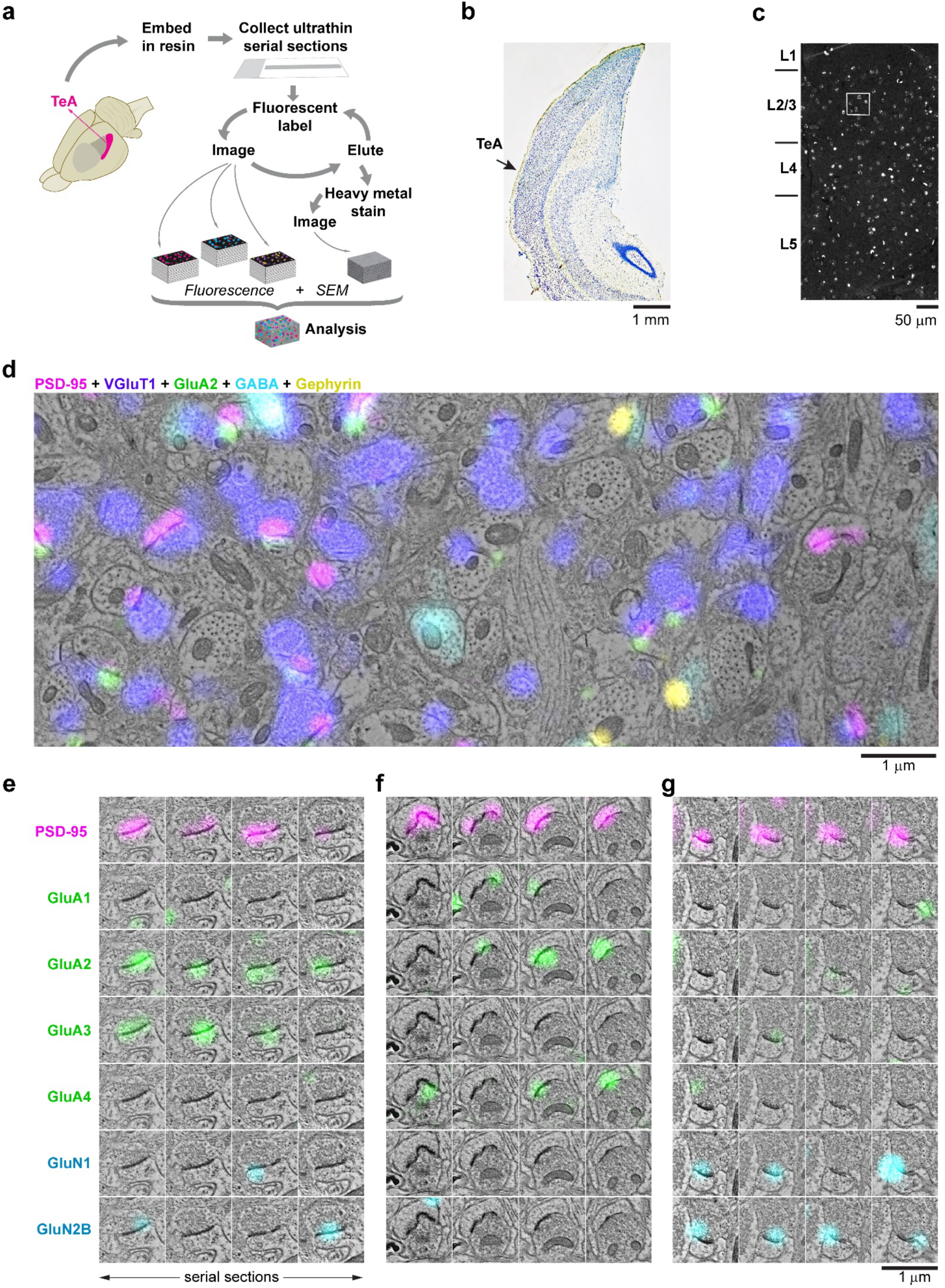
Conjugate array tomography enables multiplex immunolabeling of ultrastructurally-identified synapses. **a**. Array tomography method for serial sectioning, multi-channel IF and SEM imaging, and 3-D reconstruction of a block of layer 2/3 neocortex from the temporal association cortex (TeA) of adult mouse brain. **b**. Nissl stain of a section adjacent to the one processed for array tomography, indicating the location of TeA. **c**. DAPI fluorescence from a 70 nm ultrathin section through TeA revealing cortical layers. Region of layer 2/3 chosen for reconstruction is boxed in white. **d**. Example of conjugate IF-SEM: overview of a larger area on one section with SEM and five IF channels (PSD-95 magenta, VGluT1 purple, GluA2 green, GABA cyan, and gephyrin yellow). **e-g**. Examples of synaptograms (IF +SEM) of 3 glutamatergic synapses with different combinations of AMPA and NMDA receptor subunits. For each synaptogram, rows represent different IF channels, and columns are serial sections through the synapse.

**Table 1:** Antibodies used for Dataset 201228. (12 antibodies, 74 serial sections, 70 nm each)

| Antibody | Source | RRID | Dilution | 2° Ab (nm) |
| --- | --- | --- | --- | --- |
| **Cycle 1** |  |  |  |  |
| GluN1 | mouse, SySy 114 011 | AB_887750 | 1:500 | 594 |
| GluA1 | rabbit, Millipore #AB1504 | AB_2113602 | 1:100 | 488 |
| VGluT1 | guinea pig, Millipore AB5905 | AB_2301751 | 1:5000 | 647 |
| **Cycle 2** |  |  |  |  |
| GluA3 | mouse, Millipore MAB5416 | AB_2113897 | 1:1000 | 594 |
| GluA2 | rabbit, Abcam ab206293 | AB_2800401 | 1:50 | 488 |
| Synapsin 1/2 | chicken, SySy 106 006 | AB_2622240 | 1:100 | 647 |
| **Cycle 3** |  |  |  |  |
| GluN2B | mouse, NeuroMab N59/36 | AB_2232584 | 1:500 | 594 |
| GluA4 | rabbit, Cell Signaling 8070 | AB_10829469 | 1:50 | 488 |
| MBP | chicken, AVES MBP | AB_2313550 | 1:100 | 647 |
| **Cycle 4** |  |  |  |  |
| Gephyrin | mouse, NeuroMab L106/93 | AB_2636852 | 1:100 | 594 |
| PSD-95 | rabbit, Cell Signaling 3450 | AB_2292883 | 1:100 | 488 |
| GABA | guinea pig, Millipore AB175 | AB_2278931 | 1:5000 | 647 |

Postsynaptic receptors define the primary functional properties of glutamatergic synapses.

AMPA-type receptors are tetramers of the subunits GluA1 – 4 in various combinations, while functional NMDA-type receptors are tetramers containing the obligatory subunit GluN1 together with GluN2A and GluN2B^60–63^, in the mature neocortex. We therefore focused detection and analysis on these subunits (Extended Data Fig.1a), with the exception of GluN2A for which we were unable to identify a suitable antibody. After immunofluorescence imaging, we poststained sections with heavy metals and imaged a subregion of layer 2/3 using scanning electron microscopy (SEM) to reveal tissue ultrastructure. We then registered immunofluorescence channels with scanning electron micrographs from each section to generate volume multiplex images aligned with sub-micrometer precision. This powerful approach allows the identification of individual synapses by ultrastructure, and concomitant examination of scaffold and receptor molecules detected by immunofluorescence in those same synapses (Fig.1e-g).

Immunofluorescent labels for pre- and postsynaptic structures generally colocalized with those respective synaptic compartments in electron micrographs. Synapsin and VGluT1 colocalized with presynaptic boutons, PSD-95 colocalized with asymmetric postsynaptic densities of excitatory synapses, and gephyrin colocalized with symmetric postsynaptic densities of inhibitory synapses (Fig.1d). GABA colocalized with presynaptic boutons of inhibitory synapses, and (less brightly) with a subpopulation of postsynaptic dendrites and somata. Glutamate receptor subunits generally colocalized with PSD-95, but we also occasionally detected receptor immunolabeling outside of synapses. In some cases, extrasynaptic labeling was found in the spine apparatus, and in multivesicular bodies in dendrites (Extended Data Fig.1d), likely reflecting transport of receptors^64–66^.

By following individual synapses across multiple sequential serial sections, we found robust staining for individual synaptic markers across sections within a single synapse (Fig.1e-g). This provides strong evidence for the specificity and sensitivity of immunolabeling at single synapses. The PSD-95 label was highly consistent, overlapping with ultrastructural postsynaptic densities in most glutamatergic synapses. In contrast, there was greater heterogeneity in receptor content for AMPA and NMDA receptor subunits (Fig.1e-g), even though transcriptomic surveys suggest that they are expressed by practically all neurons in mouse TeA (Extended Data Fig.2). We therefore set out to quantify this receptor content across the synapse population.

### Proteometry of individual glutamatergic synapses reveals heterogeneity

We manually annotated 410 glutamatergic synapses based on visual inspection of electron microscopy images in the conjugate IF-SEM dataset (volume 12.1 x 12.1 x 2.6 μm^3^; Fig.2a; see Methods) and subsequently quantified immunofluorescence for each channel associated with each synapse. This revealed consistent labeling of PSD-95 and glutamate receptor antibodies within annotated synapses (Fig.2a).

**Figure 2.**
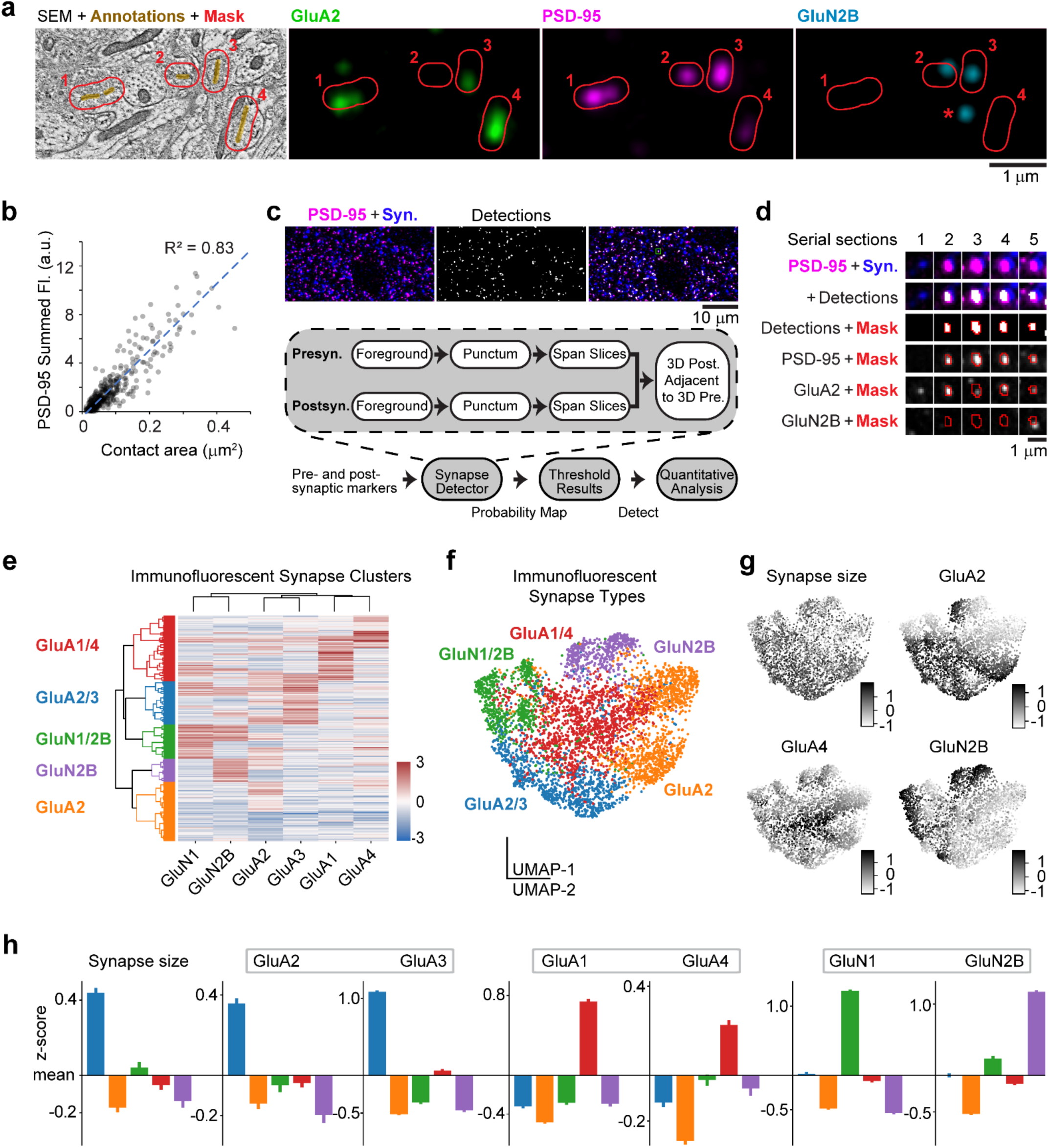
Proteometry of individual glutamatergic synapses reveals heterogeneity. **a**. Manual annotation of synapses from the IF-SEM dataset. For every synapse, immunofluorescence intensities are measured within the masked area (in red), which expands the annotation by 160 nm (see Methods). In this section, synapses 1, 3, 4 are positive for GluA2; synapses 1-4 positive for PSD-95; and synapses 2, 3 positive for GluN2B. One GluN2B-positive punctum (asterisk) is not associated with a synapse. **b**. Scatterplot reveals a strong correlation between the synapse contact area and PSD-95 immunofluorescence as quantified in the conjugate dataset (n=410 synapses). **c**. Top, example of PSD-95 and synapsin immunofluorescence used for automated synapse detection and immunofluorescence measurements at synapses in the IF-only dataset. Below, method for automated detection of synapses from immunofluorescence data using probabilistic synapse detection. **d**. Synaptogram with detections and mask. **e**. Data-driven clustering groups glutamatergic synapses from the IF-only dataset into distinct clusters based on immunolabeling characteristics (n=4,767 synapses). **f**. UMAP plot (colored by clusters defined in Panel e) shows clustering of distinct glutamatergic synapse subclasses in the IF-only dataset. **g**. Grayscale coding of UMAP projections by synapse size (summed PSD-95 immunofluorescence) and glutamatergic receptor expression in the IF-only dataset. **h**. Comparison of synaptic properties of each cluster; colors as in e and f.

Analysis of the conjugate IF-SEM dataset showed that immunofluorescence measurements correlate well with ultrastructural data. Critically, we found a strong correlation (R^2^ = 0.83) between synapse contact area (measured from SEM images) and summed PSD-95 content (measured from immunofluorescence images, Fig.2b). This provides a key validation, enabling future studies to estimate synapse size from immunofluorescence alone when using approaches such as expansion microscopy that do not support ultrastructural measurements with the same rigor as electron microscopy.

Motivated by the greater efficiency of immunofluorescence-only methods, which do not require labor-intensive electron microscopy imaging and analysis, we used the “ground truth” validation of synapses in our IF-SEM conjugate dataset to test the accuracy with which the molecular composition of synapses can be analyzed with immunofluorescence alone. To achieve unbiased automated detection of synapses in the IF-only dataset, we used a synapse detection tool developed specifically for array tomography^67^ (Fig.2c; see Methods). This tool reliably detected annotated synapses in the conjugate dataset based on co-localization of synapsin and PSD-95 (Extended Data Fig.3). To minimize false positives, we set the probability threshold at 0.75, which resulted in 13% false positives and 23% false negatives, with detection failures biased towards the smallest synapses and those with very low levels of PSD-95 immunoreactivity.

This synapse detection tool identified 4,767 glutamatergic synapses in the larger IF-only dataset (volume 46 x 44 x 3 μm^3^). We used the synapse detection area as a mask to quantify immunofluorescence for each channel within individual synapses, excluding outliers with detection area > 2.5 μm^2^ (Fig.2d, Extended Data Fig.4). The overall synapse size distribution and immunofluorescence measurements were comparable between the IF-only and conjugate dataset (Extended Data Fig.5).

Grouping synapses based on receptor content (average immunofluorescence intensity of each synapse) by Ward clustering and silhouette analysis identified five distinct clusters (Fig.2e and Extended Data Fig.6). UMAP dimensionality reduction showed a compact grouping of clusters consistent with variations in size and expression of receptor subtypes (Fig.2f).

Grayscale coding of UMAP projections by synapse size (summed PSD-95 immunofluorescence) or fluorescence intensity for individual channels revealed structured variation in molecular characteristics of single synapses (Fig.2g and Extended Data Fig.2).

The blue cluster contains large synapses with high levels of GluA2 and GluA3, and low levels of NMDA receptor subunits; while the green and purple clusters contain small synapses with low levels of GluA2 and GluA3, and high levels of GluN2B subunits; and the red cluster contains high levels of GluA1 and GluA4 (Fig.2h). The orange cluster contains medium-size synapses with high levels of GluA2 but not GluA3, as well as smaller synapses with generally low levels of any receptors. The presence of a distinct cluster positive for GluN2B but negative for GluN1 (purple) is puzzling, since GluN1 is considered an obligatory subunit for NMDAR channel function^60,68^. This result may reflect the low average number of NMDA receptors per synapse^69–72^, resulting in false negatives when immunolabeling for individual receptor subunits; indeed, for this reason many immuno-electron microscopy studies of NMDA receptor distribution at synapses have used cocktails of antibodies against several subunits^66,73^. It is possible, however, that this could also reveal a previously undescribed population of GluN1-negative NMDARs (that are presumably functionally inactive), or separate trafficking routes for the two NMDAR subunits. Similar synapse groupings are evident in the conjugate dataset where synapses are manually annotated, precluding a potential artifact introduced by the automated synapse detection (Extended Data Fig.7).

Ward clustering of the receptors yielded three groupings, identical in both the IF-only and the conjugate AT datasets: GluA2 and GluA3, GluA1 and GluA4, and GluN1 and GluN2B (Fig.2e and Extended Data Fig.7). Thus, AMPA and NMDA receptors cluster separately, and short C-tail AMPAR subunits (GluA2 and A3) cluster separately from long C-tail subunits^74^ (GluA1 and GluA4). Synapse clustering mirrors the three distinct biochemical and functional categories of receptor subunits. The overall consistency of the results using two different methods supports their validity and shows that immunofluorescence-only analysis is well suited to detect and quantify molecular properties of synapses.

### Receptor content varies with synapse size

Visual inspection of synapses from the conjugate dataset revealed that many small synapses had low levels of immunofluorescence for AMPAR subunits and PSD-95 (Fig.3a) or were completely immunonegative for those channels (Fig.3a-c), but most of them were immunopositive for NMDAR subunits^12,66,73,75^. In comparison, larger synapses had strong PSD- 95 immunolabeling in multiple sections overlapping with the ultrastructural postsynaptic density, and abundant expression of AMPAR subunits (usually GluA2 and GluA3), but much lower levels of NMDAR subunits (Fig.3d).

**Figure 3.**
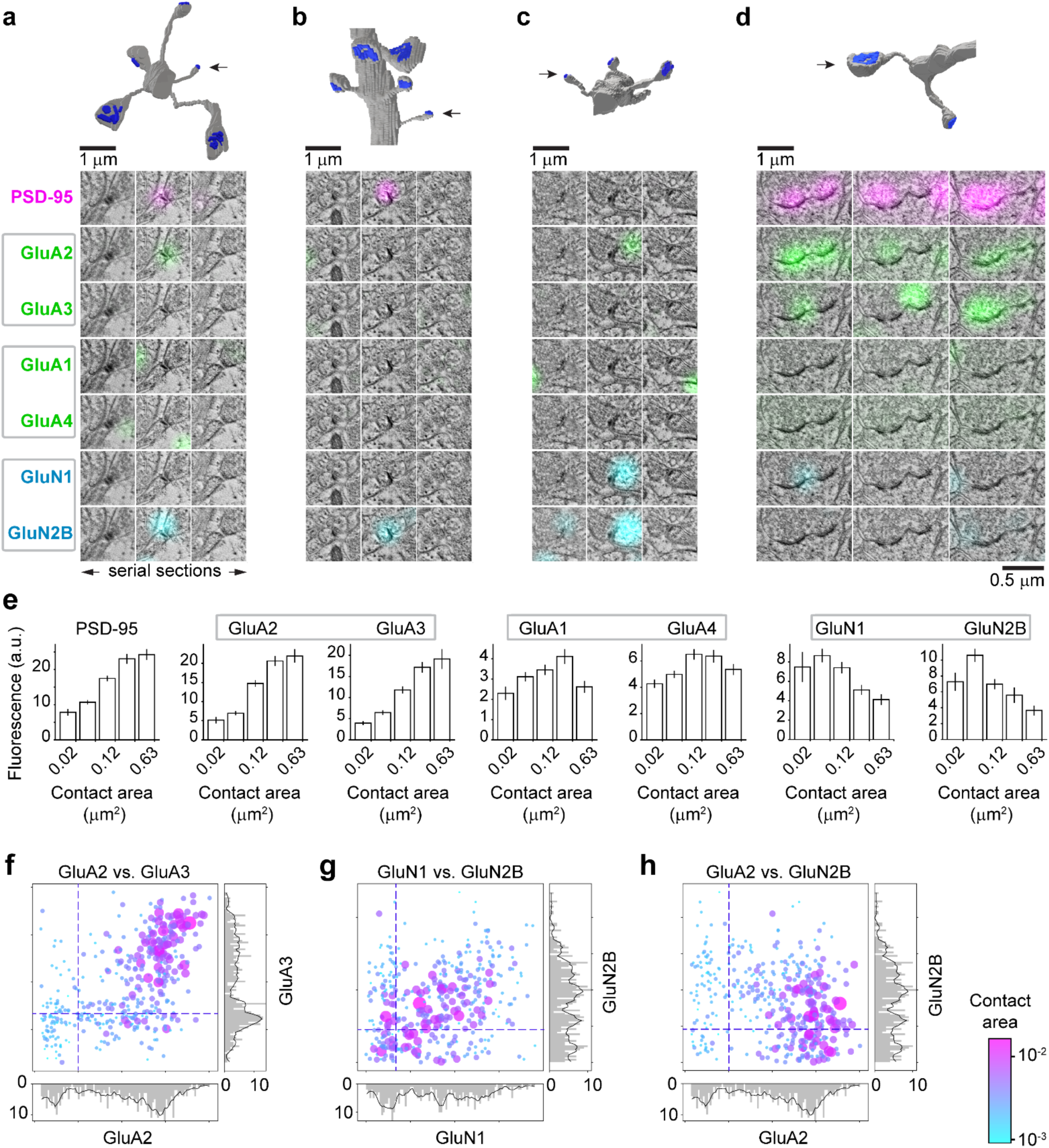
Small synapses are NMDAR-rich; large synapses are AMPAR-rich. **a-c**. Synaptograms of small synapses with a reconstruction of the parent dendrite on top. The synapse is indicated by an arrow; postsynaptic densities on spines are in blue. Fluorescence channels in this and subsequent figures are grouped following the data-driven clustering of glutamatergic receptor subunits shown in Fig.2e. **d**. Synaptogram of a large perforated synapse, showing much brighter immunolabeling for AMPARs and much weaker for NMDARs, compared to the small synapses in a-c. **e**. Histograms illustrate relationship of synapse composition to contact area in the conjugate IF-SEM dataset. X-axis histogram bins on log scale (n=410 synapses total; synapse counts per bin are n=41, n=158, n=134, n=63, and n=14 from smallest to largest contact area). **f-h**. Scatterplots comparing mean intensity across fluorescence channels at annotated synapses (conjugate IF- SEM dataset). Synapse contact area is encoded by the size and color of each dot. Dashed lines represent average fluorescence of each channel at GABAergic synapses from the same dataset, as an estimate of background.

To quantify proteometry in relation to synapse size, we measured receptor subunit immunofluorescence (average intensity per voxel) across five logarithmically-scaled synapse size bins (Fig.3e). Immunofluorescence for PSD-95 and AMPARs increased with synapse size, while NMDAR immunofluorescence declined. Among AMPAR subunits, GluA2 and GluA3 showed the strongest dependence on synapse size^12,73,75^. Functional NMDARs in adult neocortex canonically contain GluN1 subunits together with GluN2A and/or GluN2B^76^; most NMDARs in the adult mouse cortex contain GluN2B^75,77–79^. The observed decrease in GluN2B subunits at the smallest synapses might imply a potential preference for GluN2A at these synapses, although we lacked a suitable antibody to measure GluN2A.

### Single-synapse analysis reveals covariance of receptor subunits

Multiplex immunolabeling allowed us to analyze how receptor subunits covary with one another and with changes in synapse size in the conjugate dataset (Fig.3f-h and Extended Data Fig.8). We found strong positive covariance between GluA2 and GluA3, particularly in the largest synapses (upper right quadrant, Fig.3f). However, many GluA2-positive synapses were negative for GluA3 (lower right quadrant). In contrast, very few synapses were GluA2-negative and GluA3-positive (upper left quadrant), and all of these synapses were small. These results are consistent with biochemical evidence from hippocampus showing that GluA2 can exist in combination with different subunits, while GluA3 is almost exclusively found together with GluA2^80^.

GluN1 and GluN2B, when present together, covaried positively and showed an inverse correlation with synapse size (upper right quadrant, Fig.3g), consistent with the size analysis in Fig.3e. A subset of synapses that were positive for GluN1 and negative for GluN2B also showed an inverse correlation with synapse size (lower right quadrant, Fig.3g). The group of synapses that were negative for GluN1 and positive for GluN2B (upper left quadrant, Fig.3g), as in the purple cluster, must be treated with caution, as explained above.

To gain a deeper understanding of the AMPA and NMDA receptor content in relation to synapse size, we plotted GluA2 vs GluN2B. Three populations of synapses emerged, likely with distinct electrophysiological properties. The largest population comprised canonical synapses positive for both the primary AMPA (GluA2) and NMDA (GluN2B) receptor subtypes, in the upper right quadrant (Fig.3h). Within this population, larger synapses tended to have more GluA2 and less GluN2B, consistent with the general trends observed for AMPA and NMDA receptors. The lower right quadrant contained synapses positive for GluA2 and negative for GluN2B. Many synapses in this group were large, consistent with the positive correlation between AMPAR content and synapse size. These synapses may completely lack NMDARs, or may have GluN2A-containing receptors. A population of very small synapses that were positive for GluN2B but negative for GluA2 (top left quadrant, Fig.3h) represent putative silent synapses, likely present in the adult brain as well as during development^81^. Very few synapses were negative for both GluA2 and GluN2B, even though receptor content was not a criterion for synapse detection in our analysis (lower left quadrant, Fig.3h). We conclude that the presence of a postsynaptic density in adult neocortex implies functional ionotropic glutamate receptors even at the smallest synapses.

### The postsynaptic target helps to predict synaptic composition

The ultrastructural information in the conjugate IF-SEM dataset allowed us to identify postsynaptic targets of individual synapses. Glutamatergic dendrites are spiny and GABA- immunonegative, whereas GABAergic dendrites are aspiny and GABA-immunopositive.

Accordingly, to distinguish synapses onto glutamatergic dendritic shafts from those onto GABAergic dendritic shafts in our conjugate IF-SEM dataset, we first determined if the dendrites were GABA-immunopositive, and then followed the dendrites through the volume to determine if the dendrite itself was spiny.

Most synapses between glutamatergic neurons terminate onto spines, though a few terminate onto dendritic shafts. Glutamatergic synapses onto glutamatergic dendritic shafts were rare (12 out of 410 synapses, 2.9%) and tended to be small, with low levels of PSD-95 immunofluorescence (Fig.4a). Glutamatergic synapses onto GABA dendritic shafts were more common (50 out of 410 synapses, 12.2%) and comprised a distinct population exhibiting high levels of GluA1 and GluA4 immunofluorescence (Fig.4b). Synapses onto GABA-positive dendritic shafts were more densely packed along the dendrites than synapses onto non-GABA shafts (compare dendrite reconstructions in 4a and 4b), suggesting that these dendrites originated from parvalbumin interneurons^82,83^.

**Figure 4.**
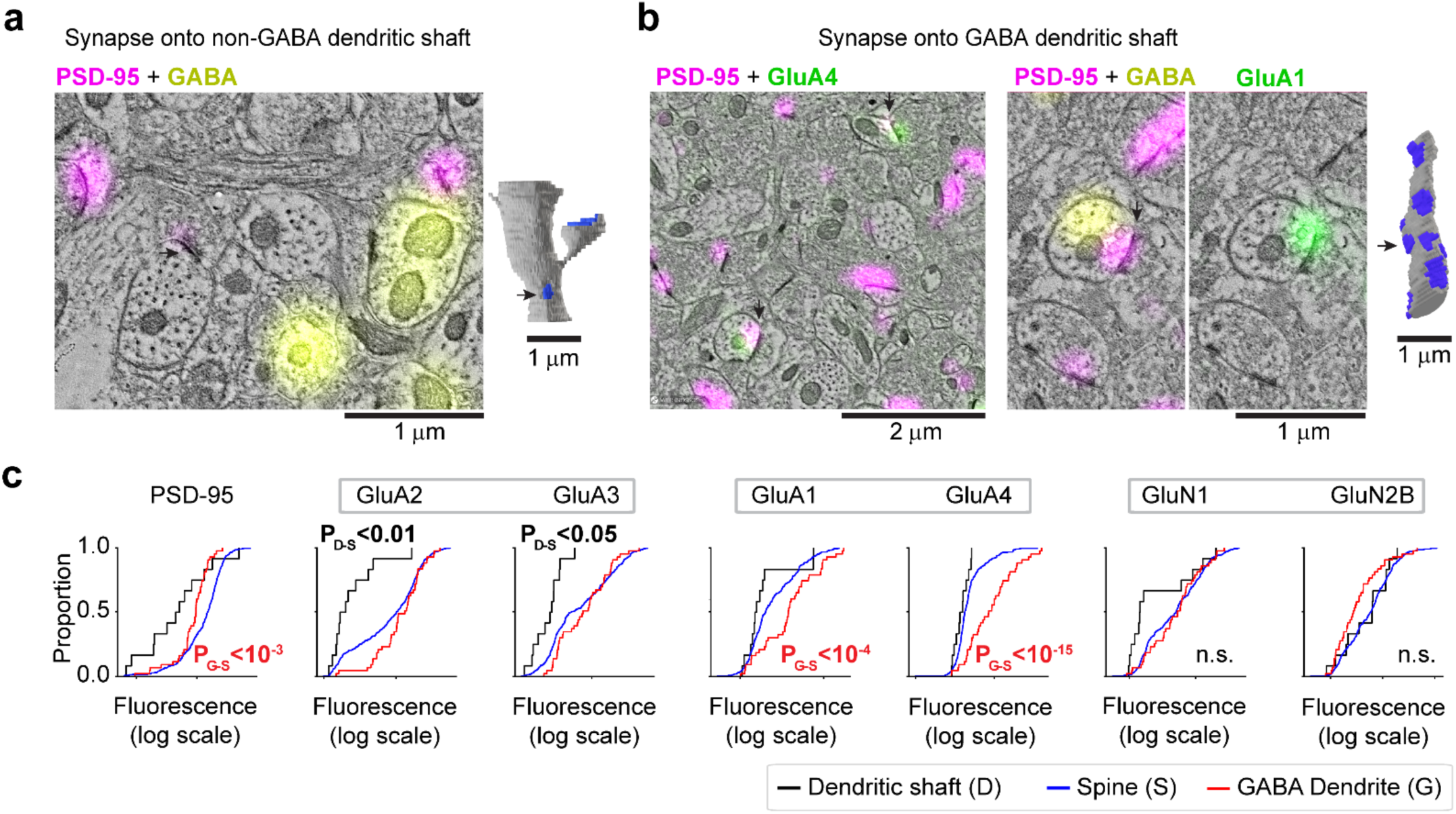
Receptor content of glutamatergic synapses depends on the postsynaptic target. **a**. Conjugate IF-SEM example of a glutamatergic synapse onto a GABA-negative dendritic shaft (reconstruction of parent dendrite on the right). These synapses tend to have weak PSD-95 fluorescence. **b**. Conjugate IF-SEM examples of synapses onto GABA-positive dendrites. The synapses on the left (arrows) are strongly immunopositive for GluA4; these synapses (higher magnification on the right) are also strongly immunopositive for GluA1. **c.** Cumulative frequency graph of receptor content of synapses, binned by postsynaptic target: synapses onto GABA dendrites have significantly more GluA1 and GluA4, and less PSD-95; synapses onto non-GABA (presumably glutamatergic neuron) dendritic shafts have significantly less GluA2 and GluA3. (n=352 spine-targeting, n=12 dendritic shaft-targeting, n=43 GABA dendrite-targeting synapses; Tukey-HSD test for all comparisons).

We quantified the receptor content across three populations of glutamatergic synapses defined by their postsynaptic target. Synapses onto GABA dendrites contained low levels of PSD-95 (Fig.4c). Synapses onto GABA-negative dendritic shafts also contained less PSD-95 than synapses onto spines, although this did not reach statistical significance, due to the small number of shaft synapses. Interestingly, synapses onto GABA-negative shafts contained less GluA2 and GluA3, while synapses onto GABA-positive shafts had significantly more GluA1 and GluA4, compared to synapses onto dendritic spines (Fig.4c). AMPARs containing GluA1 and GluA4 without GluA2 exhibit fast kinetics and high calcium permeability^84^, which position them to support high activity levels, and suggests distinct plasticity mechanisms at excitatory synapses onto inhibitory interneurons^63,84,85^.

### Immunofluorescence-only analysis at scale confirms heterogeneity of glutamatergic synapses

To test whether key features of synapse diversity can be detected using immunofluorescence array tomography, we repeated the analysis of receptor content in relation to synapse size using the larger IF-only dataset (Fig.5b). Here we used the summed PSD-95 immunofluorescence to estimate synapse size, based on the strong correlation between synapse contact area and PSD-95 immunofluorescence in the conjugate dataset analysis (Fig.2b). Results from the IF-only dataset were consistent with the conjugate dataset: GluA2 and GluA3 subunit content scaled with synapse size, while GluN1 and GluN2B content decreased.

Scatterplots of receptor co-variance were likewise consistent with the results from the conjugate dataset. GluA2 and GluA3 covaried at synapses where they are both present (upper right quadrant, Fig.5c); a group of synapses that was positive for GluA2 and negative for GluA3 was evident (lower right quadrant, Fig.5c); and a small group of synapses was negative for GluA2 and positive for GluA3 (upper left quadrant, Fig.5c). Plotting GluN1 against GluN2B (Fig.5d) showed the characteristic inverse dependence of NMDAR content on synapse size.

This plot also revealed four groupings, one in each quadrant, with different combinations of GluN1 and GluN2B, consistent with the corresponding scatterplot from the conjugate dataset (Fig.3g). The plot of GluA2 against GluN2B revealed a grouping of synapses that were positive for GluA2 and for GluN2B in the upper right quadrant, and a grouping of synapses that were negative for GluA2 and positive for GluN2B in the upper left quadrant. In both groups, the synapses with high GluN2B content were small. The synapses that were positive for GluA2 but negative for GluN2B in the lower right quadrant were large, while the synapses that were negative for GluA2 and negative for GluN2B in the lower left corner were mostly small, consistent with the conjugate data analysis. Overall, the general patterns seen in the IF-only dataset agree with those in the conjugate dataset.

To assess information about the role of the postsynaptic target neuron in the IF-only dataset, we examined GABA immunofluorescence in the volume surrounding the synapse to identify putative synapses onto GABAergic interneurons. We first validated this metric using the conjugate IF-SEM dataset, verifying that excitatory synapses in the vicinity of high GABA immunofluorescence were more likely than random synapses to terminate onto GABA targets. Indeed, when we selected the top 2% of synapses ranked by GABA immunofluorescence for ultrastructural examination, 88% of them were found to be onto GABAergic postsynaptic targets. This fraction decreased to 43% for the top 5%. In comparison, only 11.5% of all synapses in the conjugate dataset were onto GABAergic neurons.

Using this approach, we found that glutamatergic synapses adjacent to high GABA immunofluorescence (top 2%) had significantly more GluA1 and GluA4, but less PSD-95 than the remaining synapses, consistent with our results from the conjugate IF-SEM dataset (Fig.5f). We also found that synapses onto these likely GABA targets had significantly less GluA2^86^ (Fig.5f). This difference in GluA2 content was not detected in the conjugate IF-SEM dataset (Fig.4c), likely due to the smaller sample size in that dataset. Summarizing, the IF-only approach recapitulates many key results found with the conjugate dataset, but can be acquired more efficiently and therefore enables analysis of a much larger number of synapses.

Importantly, our analysis of the conjugate IF-SEM dataset validates the IF-only approach for future studies seeking to compare across multiple experimental conditions, or where equipment or expertise is not available for the more demanding conjugate IF-SEM approach.

### NMDAR content of large glutamatergic synapses is related to spine neck diameter

The large number of synapses detected in our IF-only dataset revealed an uncommon feature of glutamate receptors distribution: while NMDAR content overall is inversely correlated with synapse size (Fig.3e, Fig.5b), a subpopulation of medium and large synapses have high GluN1 and/or GluN2B immunofluorescence (Fig.5d). Deeper inspection of this population suggests that these synapses also contain high levels of GluA2 (Fig.5e). This caught our attention, because high levels of NMDAR likely endow these synapses with distinct functional properties. To examine the ultrastructure of these synapses, we leveraged the conjugate dataset where this population is also present, but less noticeable due to the smaller number of synapses (Fig.3g,h). To our surprise, many NMDAR-rich spines had a distinctive morphology, with exceptionally wide spine necks (Fig.6a).

The diameter of the spine neck may influence its function^87–91^, and the conjugate dataset allows systematic analysis of the relationship between spine neck, synapse size, and receptor content. As previously reported, spine head diameter correlated closely with synapse contact area^8^ (Fig.6b). However, we found that spine head diameter was independent of spine neck diameter (Fig.6c). We therefore partitioned large spines (head diameter ≥ 250 nm) into those with thin necks (diameter < 180 nm) and thick necks (≥180 nm). The average head diameter of these two spine populations was similar (Fig.6d), but as expected the thick-necked spines had a significantly lower ratio of head to neck diameter (Fig.6e).

Plotting GluN2B immunofluorescence against neck diameter revealed a positive correlation between neck diameter and NMDA receptor content (Fig.6f), while GluN2B showed a weak negative correlation to synapse size (Fig.6g), consistent with our synapse size analyses (Fig.3e, Fig.5b). Separate analysis of the synapses onto thin-necked versus thick-necked dendritic spines confirmed that thick-necked spines have higher GluN2B content (Fig.6h), and higher levels of GluA1, suggesting that these synapses have a potential for greater plasticity^23,74,76^.

**Figure 5.**
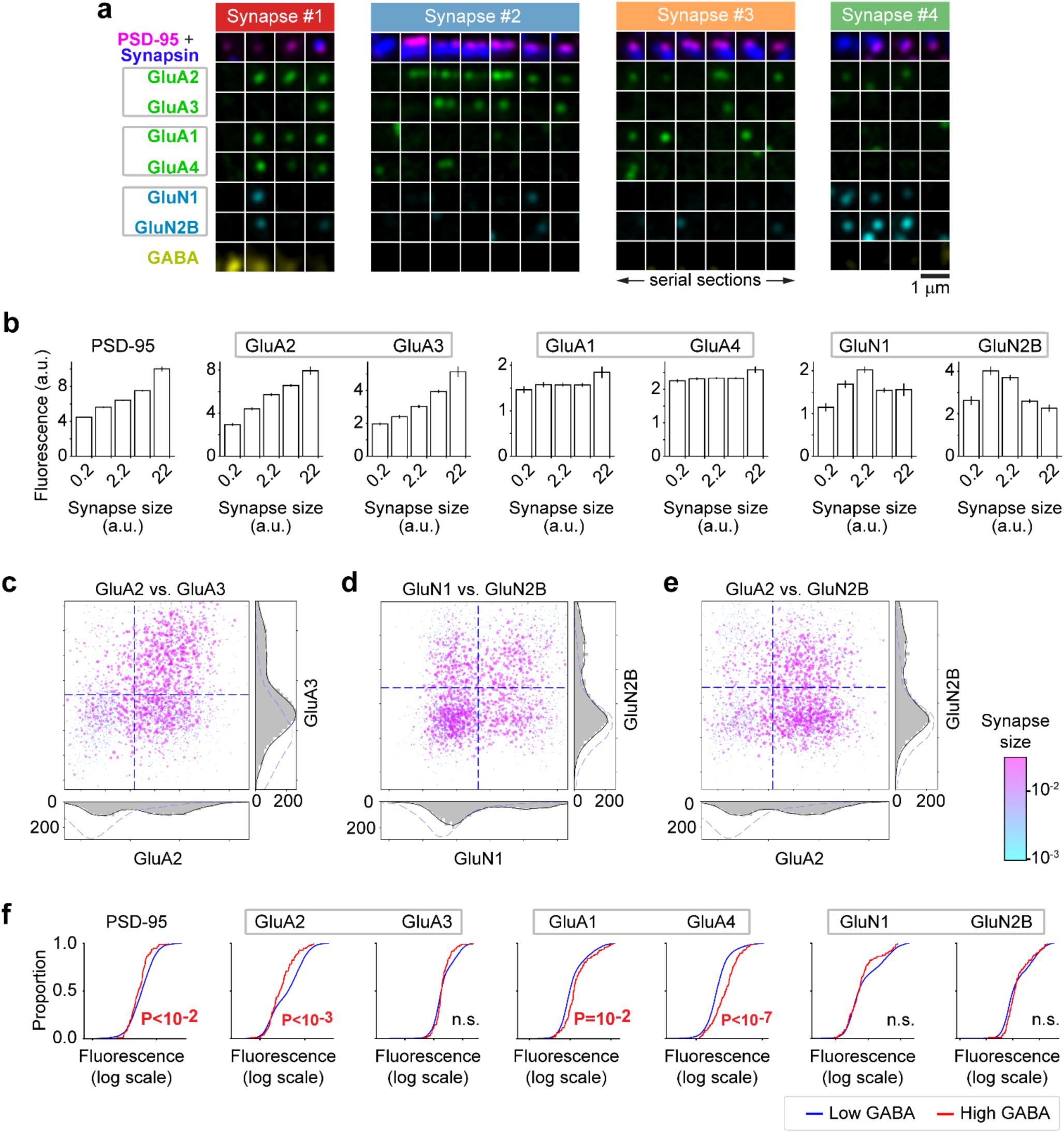
IF-only analysis at scale confirms key features of glutamatergic synapse heterogeneity. **a**. Synaptograms illustrate synapses of different sizes with different receptor composition. Synapses are color-coded at the top of each synaptogram by the corresponding synapse cluster as defined in Fig.2e,f. **b**. Histograms illustrate the relationship of synapse composition to synapse size (estimated by PSD-95 summed immunofluorescence). X-axis histogram bins on log scale (synapse counts per bin are n=583, n=1016, n=1646, n=1355, and n=134 from smallest to largest). **c-e**. Scatterplots comparing mean intensity across fluorescence channels at automatically detected synapses (IF-only dataset). Synapse detection volume is encoded by the size and color of each dot. Histograms show distribution of intensities for each axis (gray) and distribution of intensities in a shuffled set of synapse puncta locations from that channel (dashed), as an estimate of background. Dashed lines on the scatter plot represent the mean value + 1 standard deviation of the shuffled synapse puncta locations from the same dataset. **f**. For the IF-only dataset, fluorescence in a dilated GABA channel (mask is detection pixels dilated by 2 pixels) is used to identify synapses onto GABA targets. Cumulative frequency graph of receptor content of synapses onto putative GABAergic dendrites vs. the rest; like the conjugate dataset, synapses onto potential GABA targets (exhibiting the strongest 2% of GABA immunofluorescence) have significantly more GluA1, GluA4 and less PSD-95. Synapses with high GABA content also have significantly less GluA2. (n=96 high GABA content synapses; n=4626 low GABA content synapses).

### Receptor content correlates more strongly with synapse ultrastructure than with parent neuron identity

The molecular composition of synapses is determined, at least in part, by the identity of the parent neurons. Accordingly, we detected differences between the receptor content of synapses onto glutamatergic vs. GABAergic dendrites (Fig.4). To further explore the influence of the parent neurons, we identified synapses within the conjugate dataset that share the same postsynaptic dendrite (Fig.7a), the same presynaptic axon, or both. As expected, pairs of synapses sharing the same postsynaptic dendrite (n=412 pairs) were more similar in receptor content than all synapse pairs, however, the magnitude of this effect was surprisingly modest.

Conversely, pairs of synapses onto a glutamatergic and a GABAergic dendrite (GABA-Glut Mix, n=15,953, Fig.7c,d) had higher divergence in receptor content than the population of all synapse pairs. To permit a more extensive visual examination of these results, we color-coded subsets of synapses based on parent dendrite identity within the UMAP projections (Fig.7b), but failed to find any clear evidence that synapses that shared the same postsynaptic dendrite grouped together. We also identified a small number of synapses sharing the same presynaptic axon (n=70 pairs), but for this limited sample we found no significant differences in similarity compared to all synapses.

Prompted by the correlation between receptor content, synapse size and spine neck size (Fig.6), we next defined three subsets of axospinous synapses based on distinctive morphology: “classical” spines with well-defined spine head and spine apparatus, including a perforated postsynaptic density and presence of presynaptic mitochondria (n=33); thick-necked spines, with a neck diameter ≥ 180 nm and head diameter ≥ 250 nm (n=22); and small spines with a head diameter < 250 nm (n=23). There was partial overlap within the first two categories (8 spines) (Fig.7e,f).

**Figure 6.**
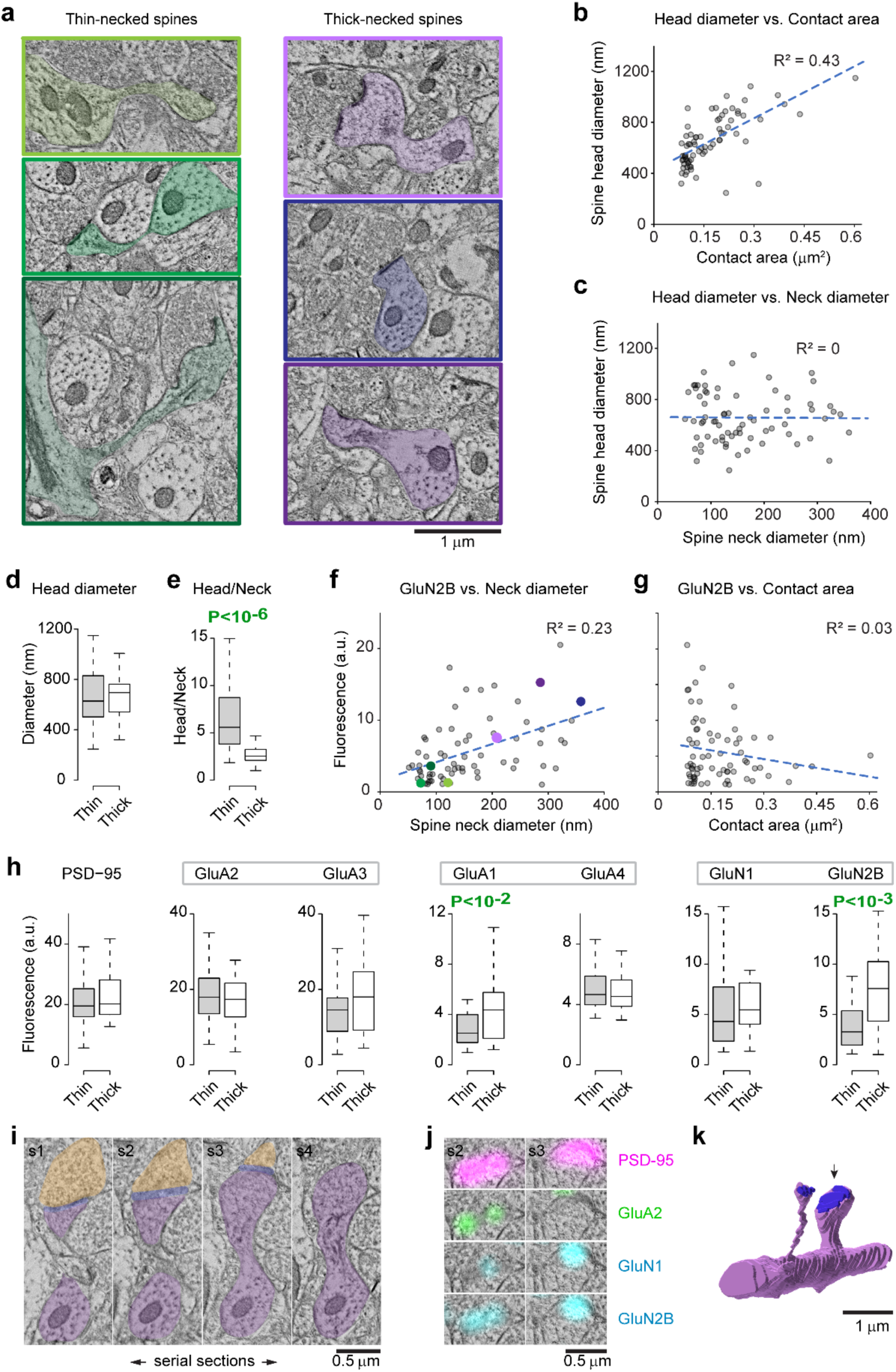
Receptor content of large glutamatergic synapses correlates with spine shape. **a**. Examples of large spines with thin necks (<180 nm diameter) vs. those with thick necks (≧180 nm diameter), with the color corresponding to the color dots in f. **b, c**. Scatterplots show that the spine head diameter has a strong positive correlation with synapse size (b), but no correlation with neck diameter (c). Only large spines (head diameter ≧ 250 nm) are included in these scatterplots, n= 72. **d, e**. Thin- and thick-necked spines have similar head diameters (d), but a significantly different ratio of head/neck diameters (e) based on a sample of 50 thin-necked spines and 22 thick-necked spines. Statistical significance (t-test) is indicated on the plot. **f**. Scatterplot shows a positive correlation between GluN2B average fluorescence and spine neck diameter of large spines. GluN2B and spine neck size are positively correlated. The colored dots correspond to the spines in c with the same color box. N=72 large spines. **g**. Scatterplot of GluN2B average fluorescence vs. contact area, showing weak negative correlation. **h**. Comparison of receptor content of thin-necked vs. thick–necked spines. Statistical significance (t-test) is indicated on the plot. Synapses onto spines with thick necks have significantly more GluA1 and GluN2B immunofluorescence. **i**. Four serial sections through a thick-necked spine. The spine is segmented out in purple, the presynaptic bouton in yellow, and the postsynaptic density in blue. **j**. Immunofluorescence for PSD-95, GluA2, GluN1 and GluN2B in two of the sections through the synapse. **k**. Volume reconstruction of the same thick-necked spine, indicated with an arrow.

Notably, we found that synapses onto ultrastructurally-defined spine types clustered together on the UMAP projection (Fig.7f) and exhibited greater similarity in receptor content than synapses sharing pre- or postsynaptic parent neurons. Synapses onto classical and thick- necked spines clustered together with other large synapses with high AMPAR content, but the two populations showed some separation (Fig.7f,g). Synapses onto small spines clustered separately with other small NMDAR-rich synapses. Synapses belonging to each of these ultrastructurally-defined spine types were more significantly similar to one another than to the population of all synapses.

We conclude that parent neuron identities have relatively little influence, and that spine morphology is a far better predictor of glutamatergic receptor content at axospinous synapses. The strong similarity of synapses onto classical spines likely reflects stability of these synapses^92–94^ and an optimization of receptor content for their primary role in robust transmission of information. This important result is consistent with a key role for synapse subclasses as distinct computational units that are not simply defined by parent neurons^95,96^.

## Discussion

Besides serving as key nodes of communication between neurons, synapses also act as independent computational units. Insight into the functional specialization of synapses in the mammalian brain is constrained by the techniques available to assess their molecular diversity. These techniques must contend with the small size and high density of synapses within the brain. Consequently, far less is known about their heterogeneity and specialization than larger-scale circuit components such as cell types and microcircuits.

Here we leverage the unique capacity of conjugate array tomography to probe the chemical architecture of synapses within the ultrastructural context of the brain. Using machine-learning classification approaches, we find that glutamatergic synapses in the neocortex group into distinct clusters based on their content of receptor subunits. Previous studies have clustered synapses based on non-receptor proteins in dissociated hippocampal cultures^97–99^, or on optically-measured colocalization of two postsynaptic scaffolding proteins^32,100^. Proteomic analyses of bulk synaptosome preparations reveal co-regulation of groups of proteins that may correlate with synapse subclasses^35^ and molecular specialization of synapses from genetically defined synapse types^101^. However, no such proteometric clustering in the intact brain has previously been reported.

Our focus on synaptic receptors permits functionally-relevant insight into the logic underlying the specialization of glutamatergic synapses. By registering volume electron microscopy with immunofluorescence, we demonstrate that specific proteomic features are associated with distinct ultrastructural characteristics, and we identify a distinct group of NMDA-rich synapses onto large thick-necked spines. Synapse diversity has been correlated with the identity of the pre- or post-synaptic parent neuron^8,102–104^, but we find that the identity of the parent neurons accounts for only a modest fraction of the molecular and ultrastructural diversity of synapses, suggesting that synapse subclass itself may be a fundamental element of neuronal circuit structure^96^.

### Limitations of the current study

Our data are from the TeA cytoarchitectonic field of a 16-month old mouse (roughly corresponding to a 50 year-old human). By avoiding inter-animal variability, our deep analysis of a single animal limits technical measurement noise, as in previous labor-intensive connectomic datasets^105,106^. TeA is homologous to human temporal association cortex, facilitating future comparison with neurosurgically resected tissue blocks (whose tissue quality is far superior to post-mortem samples)^107–111^, since these blocks are typically resected from the middle temporal gyrus to access an underlying epileptic focus^107^. The manual annotation of synapses in the electron microscopy images limited the size of the conjugate array tomography dataset compared to the more efficient, automated synapse detection used for IF-only analysis. Nevertheless, the conjugate array tomography dataset allowed rigorous validation of the larger IF-only dataset.

We used a carefully selected and validated set of commercial antibodies to assess the distribution of receptor subunits in a reproducible manner^10,112–114^. However, the performance of the antibodies was variable. GluA1 showed less sensitivity and higher background than the other antibodies, but we retained GluA1 in our dataset based on its colocalization with PSD-95, and robust labeling of select individual synapses (e.g. Fig.1f and Fig.5a). We detected a cluster of synapses immunopositive for GluN2B but immunonegative for GluN1, even though GluN1 is an obligatory subunit for functional NMDA receptors^60^. This observation must be treated cautiously because it could arise from stochastic noise, reflecting limited labeling efficiency and the fact that there are fewer copies of NMDA receptors per synapse compared to AMPA receptors^69–72^. However, it could also suggest a previously undescribed population of functionally inactive GluN1-negative NMDARs, or separate trafficking routes for the two subunits.

Our automated synapse detector enabled efficient analysis of synaptic populations using immunofluorescence only. This approach is more accessible and scalable than conjugate IF- SEM array tomography. Furthermore, the validity of the IF-only approach was largely supported by direct comparison of the immunofluorescence and electron microscopic detection of synapses. However, automated synapse detection based on immunofluorescence failed to reliably identify the smallest synapses, which span only one or two sections and have low levels of immunoreactivity for synaptic markers.

### Factors predicting receptor expression

We found that glutamatergic synapses in the TeA group into subclasses based on glutamate receptor content, implying distinct physiological roles. For example, the blue cluster of large synapses with high expression of GluA2/GluA3 and low expression NMDAR subunits represents strong synapses, while the purple and green clusters of small synapses low in AMPARs and high in NMDARs likely include weak or silent synapses. The discrete divisions between groupings of synapses that we observe based on receptor content are reminiscent of the bimodal distributions of synapse sizes that have been reported between layer 2/3 pyramidal neurons in mouse neocortex^115^. Extending this approach to analyze more synapses in a larger neocortical dataset through array tomography or multiplex expansion microscopy^99,117–120^ would likely reveal additional uncommon clusters and would also allow clear identification of subgroups within the clusters reported here. Likewise, extending this analysis to detection of additional synaptic proteins and features, and to examination of additional brain structures, species, and stages of neurodevelopment or aging would likely uncover new discrete clusters of synapse subclasses, as well as continuous sources of variation within existing subclasses.

Many factors contribute to the diversity of glutamatergic synapses, including the phenotype of the parent pre- and postsynaptic neurons, the regional neurochemical milieu of the synapse, and the history of neural activity. Our data confirm that the postsynaptic target is an important predictor of the receptor content of excitatory synapses. We found markedly lower levels of PSD-95, GluA2, and GluA3 at glutamatergic synapses onto excitatory dendritic shafts, compared to those onto dendritic spines. We also detected differences in the receptor composition of synapses onto GABAergic dendrites relative to those onto glutamatergic dendrites. Although our data did not allow clear identification of different subclasses of GABAergic interneurons, synapses onto GABAergic dendritic shafts expressed high levels of GluA1 and GluA4. Based on the fast kinetics and high calcium permeability of these receptor subtypes^84^, this receptor specialization is consistent with suggestions that inhibitory neurons may play a special role in high-frequency signaling.

### Spine morphology and receptor content

Synapses onto large “classical” mushroom-shaped spines contained high levels of AMPARs and few NMDARs, whereas those onto small spines contained relatively fewer AMPARs and more NMDARs, consistent with previous reports^12,73,75^. Notably, we also found that large spines with thick necks comprised a distinct group associated with both high AMPAR and high NMDAR content. The existence of specialized thick-necked spines has been suspected based on the skewed distribution of neck sizes^121^, but there was no previous evidence that such spines have a distinct molecular composition.

Spines with thick necks permit rapid diffusion of calcium and other signaling molecules into the dendrite^88,89^, where they could engage signaling pathways not localized to the spine, allowing thick-necked spines to exert more influence over the parent dendrite. This is consistent with their high levels of NMDAR channels, whose slow kinetics and high calcium permeability play a special role in synaptic plasticity, and probably also in learning and memory. Conversely, a thicker neck would also support greater diffusion from the parent dendrite into the spine, making these spines more sensitive to cell- and dendrite-wide signaling and allowing more rapid changes to the chemical composition of the synapse^88,89,122^. Restricted spine neck size may also reduce electrical conductance, providing electrical isolation for thin-necked but not thick-necked spines^90,91,123^, although this idea remains controversial due to difficulty of measuring spine voltage^88,89,122,124^.

### Histological identification of putative silent synapses

We identified a subgroup of small synapses that contained NMDARs but no detectible AMPARs. These synapses likely correspond to electrophysiologically-defined postsynaptically- “silent” synapses, which have been posited as a key players in long-term potentiation^125–127^.

Because immunocytochemistry detects protein directly, our study gives independent orthogonal support for the reality of “silent” synapses. It remains possible that some of these synapses may actually express AMPARs at levels below the threshold for antibody detection; whether so few AMPARs would be sufficient to remove the voltage-dependent block of NMDARs is debatable. While previous work described synapses lacking NMDARs in juvenile rodents^12,73,75^, our study supports the notion that silent synapses persist through adulthood and are thus likely to continue to be involved in circuit remodeling, building on a recent study of younger adults^81^.

### Synapse subclasses as computational units

We found that AMPA and NMDA receptor content of axospinous synapses were only weakly predicted by pre- or postsynaptic parent neuron identity. This is surprising; the molecular composition of a synapse is surely constrained by the pre- and postsynaptic neurons’ transcriptomes, and synapses that share the same parent neurons tend to be more similar in size than random synapses^102,104,128^. However, we find that ultrastructural features are much stronger predictors of glutamate receptor content. For example, synapses onto anatomically defined “classical” spines with a large mushroom head showed much greater molecular similarity to other synapses onto other classical spines than to synapses sharing the same pre- or post-synaptic parent neuron (Fig.7). These results are consistent with a key role for synapse subclasses as distinct computational units that are not simply defined by parent neurons^95,96^.

**Figure 7.**
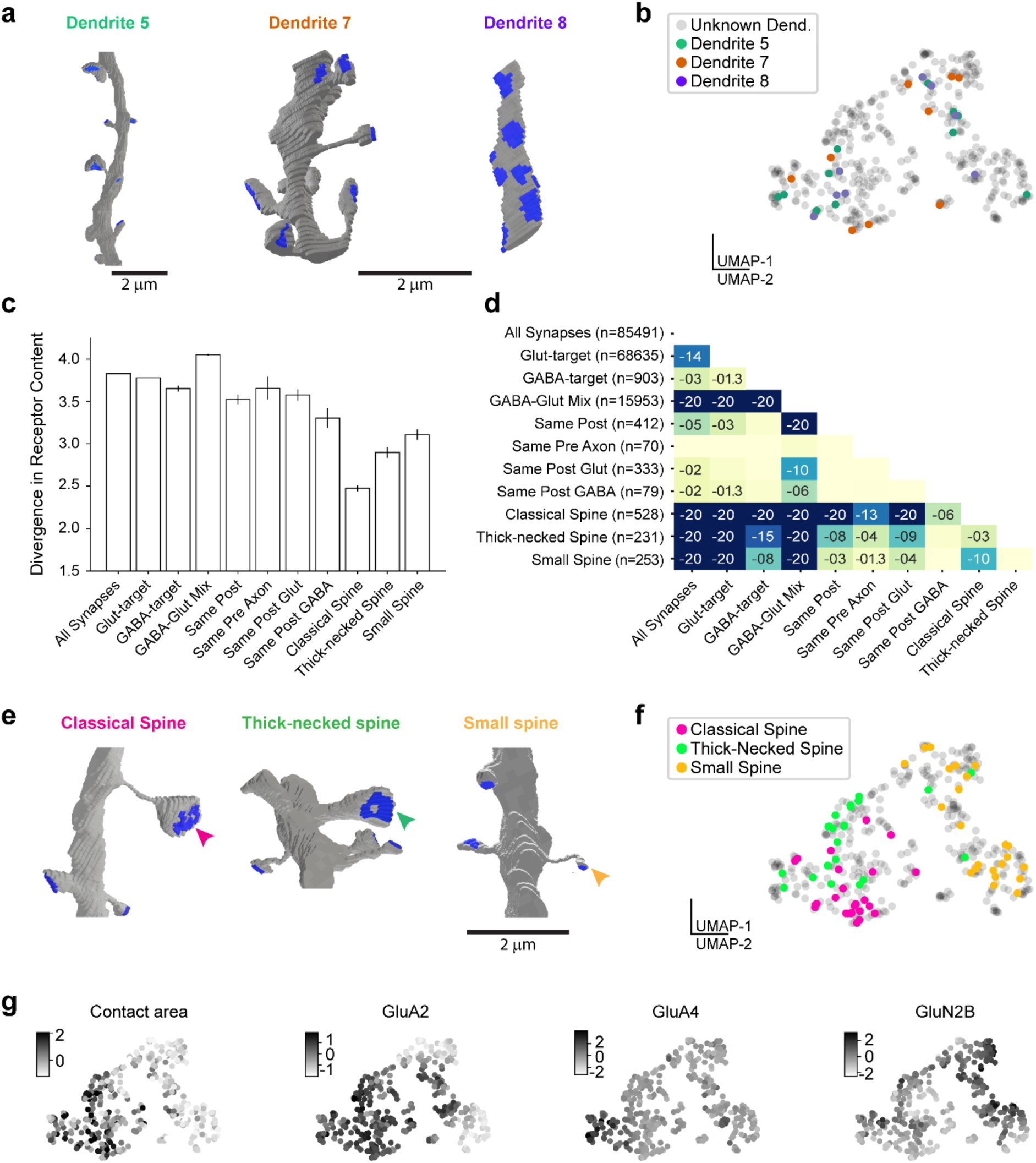
Receptor content of synapses is only partially determined by the pre- and postsynaptic neuron. **a.** Example of synapses onto the same dendrite, with postsynaptic densities in blue. Dendrite #8 is GABA-immunopositive. **b**. UMAP plot with synapses targeting the same dendrite identified by the same color. Synapses onto the same postsynaptic target show only weak evidence of clustering. **c**. Analysis of the divergence in receptor content of synapses, depending on the postsynaptic target, presynaptic axon, and spine morphology. Synapses sharing the same postsynaptic dendrite are more similar in receptor content compared to all synapses, but the difference is very small. In comparison, synapses onto morphologically-defined spine types show much higher similarity. (All synapses n=85,491 pairs; glutamate-targeting n=68,635 pairs; GABA-targeting n=903 pairs; GABA- and Glut-targeting mix=15,953 pairs; same postsynaptic dendrite n=412 pairs; same presynaptic axon n=70 pairs; same postsynaptic glutamate-targeting n=333 pairs; same postsynaptic GABA-targeting n=79 pairs; classical spine n=528 pairs; thick-necked spine n=231 pairs; small spine n=253 pairs) **d**. Table with all the statistics for c, numbers represent the exponent of the probability as estimated by Tukey HSD test. **e**. Examples of morphological groups of dendritic spines: “Classical” spines contain a spine apparatus postsynaptically, a mitochondrion presynaptically, and a perforated postsynaptic density. Thick-necked spines are spines with a neck diameter ≥ 180 nm. Small spines are defined as spines with a head diameter < 250 nm. **f**. UMAP plot showing clustering of synapses onto the three different morphological groups of spines. **g**. Grayscale coding of UMAP projections by synapse size (contact area) and glutamatergic receptor expression in the conjugate IF-SEM dataset.

### Implications for future research

The molecular composition of synapses changes dramatically during postnatal development. Synapses are especially modifiable before maturity, but remain highly plastic even in the adult mammal. Multiple factors, including the tissue environment and the location on the postsynaptic neuron, can influence the morphology and composition of synapses. The precise history of activity at each synapse and the resulting pattern of exposure to diffusible neuromodulators can likewise drive synaptic variability. However, as sites of memory storage, synapses must also possess a capacity for “stability”. Physiological literature has implicated transitions between discrete states as a primary basis for glutamatergic synaptic plasticity^22,23,129–135^, and theoretical studies propose discrete states with barriers to transition as a basis for synaptic stability^130,136–139^. This raises the possibility that our subclasses from a single proteometric snapshot may represent samples from a dynamic population, where activity-dependent plasticity is driving transitions by individual synapses over energy barriers separating discrete stable states. New avenues for imaging glutamate receptors in behaving animals will be instrumental in addressing these questions^140^.

Recent technical improvements are driving rapid advances in connectomics based on volume electron microscopy^105,106,141–146^. However, the functional impact of connections between two neurons or brain structures depends on the strength and physiological features of each synapse in that connectome. Improved methods for qualitative and quantitative estimation of these features from ultrastructural evidence will be essential for future generations of connectomics. Ultrastructural correlates that distinguish excitatory from inhibitory synapses were identified 50 years ago^147–149^, while more recent work has begun to elucidate the relationship between synapse size and synaptic strength^1,2,4,116,150,151^ or even ultrastructure and neurotransmitter content^152^. Conversely, new methods in expansion microscopy and multiplex super-resolution imaging can provide increasingly rich molecular information about single synapses^97,103,117–120^, although at the expense of ultrastructural context (but see^153–155^). Our direct measurement of receptor content rigorously aligned with ultrastructural features at single-synapse resolution therefore represents an important step toward a functional interpretation of connectomes.

Animal models and genetic studies of debilitating neuropsychiatric disorders, including obsessive-compulsive disorder, autism, and schizophrenia, point to synaptic proteins as a critical locus of the underlying circuit dysfunction^37–42,156–158^. While numerous studies focus on identifying the cell type or brain region that underlies these disorders, our results suggest that synapse subclass may be an important but overlooked organizational principle for the etiologic mechanisms and treatment of these disorders^98^. Advances in array tomography, expansion microscopy, super-resolution imaging, and quantitative proteomics are ideally positioned to offer new insights.

## Acknowledgments

We thank Forrest Collman, Uygar Sümbül, Michael Hawrylycz and members of the Owen lab for helpful comments. We also wish to thank Allen Institute founder, the late Paul G. Allen, for his vision, encouragement, and generous support.

## Author Contributions

Conceptualization: KDM, SJS, RJW; Methodology: KDM, AKS, JS, SJS, RDW, SFO; Software: AKS, SFO; Validation: KDM, AKS, JS, RJW, SFO; Formal Analysis: KDM, AKS, JS, SJS, RJW, SFO; Investigation: KDM, AKS, JS, SJS, RJW, SFO; Resources: SJS, RJW, SFO; Data Curation: KDM, AKS, JS, SJS, RJW, SFO; Writing: KDM, SJS, RJW, SFO; Supervision: KDM, SJS, RJW, SFO; Funding Acquisition: KDM, SJS, RJW, SFO.

## Methods

Specimen preparation and array tomography methods closely parallel those previously described in detail^10,43,44,67,113^. Those details are summarized briefly here, alongside new particulars where needed.

### Animals

Adult C57BL/6J mice were used for this study. The conjugate volume and the larger IF-only volume are from a 16-month old male mouse (M4926). All animal procedures were performed in accordance with the University of North Carolina animal use committee’s regulations.

Mice were deeply anesthetized with pentobarbital/ketamine; after a brief saline flush, they were perfusion-fixed with a mixture of 2% glutaraldehyde/2% formaldehyde in 0.1 M cacodylate buffer, pH 7.0, with the addition of 2 mM CaCl2 and 2 mM MgSO4. The brain was removed and postfixed overnight at 4°C in the same fixative. 50 μm-thick sections were cut on a Vibratome and collected in 0.1 M phosphate buffered saline. Relevant sections were cryoprotected in 30% glycerol and quick-frozen onto an aluminium block cooled on dry ice, to aid in reagent penetration. Cold sections were quickly placed into vials containing ethanol + 0.1% uranyl acetate at -70° in a Leica AFS machine. After 24 hours dehydration, ethanol was progressively replaced with HM-20 Lowicryl resin at -50° C. After plastic infiltration was complete, sections were sandwiched between sheets of ACLAR polychloro-trifluroethylene plastic, and exposed to long-wavelength UV light as incubation temperature was gradually increased over 18 h to room temperature. The ACLAR was peeled off the polymerized wafers and small blocks containing TeA auditory association cortex were cut out of the sections and glued to polymerized cylinders of epoxy plastic. The blocks were mailed to Stanford University for subsequent thin sectioning, immunolabeling and fluorescence imaging.

### Substrate coating

Schott Nexterion A+ coverslips (Fisher Scientific) were used as a substrate for collection. The functionalized aminosilane glass was carbon coated using a Cressington 308R Coater to ensure strong adhesion of the section ribbon to withstand subsequent harsh treatments, and to improve conductivity for the scanning electron microscope. Coverslips were coated three times for 9.9 seconds each, resulting in a total deposit of ∼ 3 nm of carbon on the surface.

### Ultrathin sectioning

The blocks were trimmed around the tissue to the shape of a trapezoid approximately 1.5 mm wide and 0.5 mm high, and glue (Weldwood Contact Cement diluted with xylene) was applied with a thin paint brush to the leading and trailing edges of the block pyramid. Ribbons of 70-nm-thick serial sections were cut on an ultramicrotome (Leica Ultracut EM UC6) and mounted onto carbon-coated aminosilane coverslips prepared as described above. After the sections dried, they were kept on a slide warmer set at 55° C for 30 minutes, and then stored until immunolabeling within 1 week from sectioning.

### Immunofluorescence labeling

The sections mounted on the coverslips were immunolabeled as previously described^10^. Sections were pretreated with freshly made sodium borohydride [1% in Tris-buffered saline (TBS), pH 7.6 for 3 min] to reduce non-specific staining and autofluorescence. After a 20 min wash with TBS, the sections were incubated in 50 mM glycine in TBS for 5 min, followed by blocking solution (0.05% Tween 20 and 0.1% BSA in TBS) for 5 min. The primary antibodies (listed in Table 1) were diluted in blocking solution and applied overnight at 4°C. After a 15-min wash in TBS, the sections were incubated with highly cross-adsorbed Alexa Fluor conjugated secondary antibodies (ThermoFisher Scientific), diluted 1:150 in blocking solution for 30 min at room temperature. Finally, sections were washed with TBS for 15 min, rinsed with distilled water, and mounted on glass slides using SlowFade Diamond Antifade Mountant with DAPI (ThermoFisher Scientific S36964). After sections were imaged, the antibodies were eluted using a solution of 0.2 M NaOH and 0.02% SDS for 20 minutes, and new antibodies were reapplied, as needed. All staining rounds included the DAPI label to facilitate image registration between rounds.

### Antibodies

All antibodies were from commercial sources and have been extensively validated for AT in previous studies^10,43,113,114,159,160^.

### Fluorescence imaging and image processing

The immunostained ribbon of sections was imaged on an automated epifluorescence microscope (Zeiss AxioImager Z1) using a 63x Plan-Apochromat 1.4 NA oil objective. To define the position list for automated imaging, a custom Python-based graphical user interface, MosaicPlanner (Dr. Forrest Collman^161^, obtained from https://code.google.com/archive/p/smithlabsoftware/), was used to find corresponding locations across the serial sections. An area of 46 x 44 μm in layers 2/3 was selected for analysis. The images were registered between staining cycles and aligned based on the DAPI signal, using the MultiStackReg plugin in FIJI^162^. Background was subtracted using the Subtract Background function in FIJI, with a rolling ball radius of 50. Images were normalized with the Enhance Contrast function in Fiji by setting saturated pixels value at 0.01%. Occasional brightly fluorescent specks of contamination were masked out to not interfere with the normalization.

This volume is referred to as IF-only volume in the Results section.

### Post-staining

Following 4 rounds of multiplex fluorescence imaging, the ribbons were rinsed with water, dried and mailed to the Allen Institute for SEM imaging. To increase contrast, the sections were first poststained for 1 min with a freshly-made solution of 0.1% KMnO4 dissolved in 0.1 N H2SO4, followed by 5% aqueous uranyl acetate for 30 min, rinsed in water, and then 1% Reynolds’ lead citrate, freshly prepared and filtered, for 1 min^43^. Sections were then rinsed in water and air dried.

### SEM imaging

A Zeiss Gemini 500 using Atlas V software was used to acquire the correlative EM dataset. A 4x4 mosaic with a size of 49.2 μm x 49.2 μm in layer 2/3 was acquired for each section, mapped with ATLAS V, with a resolution of 3 nm/px. This was done with a stage bias energy of 5 kV, allowing for a landing energy of 1.75 keV, and at a working distance of 3.25 mm.

### Stitching & registration

Stitching and registration were modelled after published methods^43^, employing render (https://github.com/saalfeldlab/render) to store transformations of individual images, and custom python code (https://github.com/AllenInstitute/render-python-apps/blob/master/renderapps/cross_modal_registration) that helped create TrakEM2 projects for each section to be set up. Within the projects, each section was stitched, roughly registered for each multiplexed section, then hand-registered using correspondences between the PSD-95 imagery and synapses seen in the SEM channel. After registration, upsampled IF and EM imagery were exported from TrakEM2 and saved as individual TIFF files. A smaller volume (ROI1: 12.1 x 12.1 x 2.6 μm) was cut out and aligned across sections with the MultiStackReg plugin, (Brad Busse) in FIJI^127^, using the SEM channel. The resulting transformations were then applied to the other channels. This small volume including the SEM channel and all the IF channels was uploaded to webKnossos^163^ to be analyzed as a conjugate IF-SEM volume.

### Image processing of the conjugate IF-SEM dataset

For the fluorescence channels, background was subtracted using the Subtract Background function in FIJI, with a rolling ball radius of 500. Images were normalized using the Enhance Contrast function, setting the saturated pixels at 0.01%. The SEM images were adjusted in FIJI by converting to 32 bit and applying an Unsharp mask (radius 2, mask weight 0.7). Contrast was enhanced using the Enhance Contrast function with saturated pixels at 0.02%, followed by gamma adjustment to 1.2, and Unsharp mask (radius 1, mask weight 0.4). The processed images were finally reverted back to 8 bit. These image processing steps were applied before further analysis of the conjugate dataset.

### Conjugate dataset annotation

Excitatory synapses were annotated in webKnossos using the “Segments tool”. Synapses were identified by ultrastructural features, such as the accumulation of presynaptic vesicles apposed to a postsynaptic thickening. In cases of uncertainty, immunofluorescence for PSD-95 was used to assist the identification of *en face* synapses, and immunofluorescence for GABA to distinguish between excitatory and inhibitory synapses. An 80-pixel brush (corresponding to 27 nm) was used to paint over the postsynaptic density of each synapse in the dataset as seen on the serial sections through that synapse. The annotation of each synapse was allocated a separate ID number as an individual segment.

### Automated detection of glutamatergic synapses

Glutamatergic synapses were detected in fully aligned and registered multichannel AT volume images using previously described synapse detection software^67,160^. Though PSD-95 is thought to be present at all, or nearly all, glutamatergic synapses^164^, it is also detectable at many obviously non-synaptic sites. PSD-95 can therefore be considered necessary but not sufficient when identifying synapses in our multiplex images. We therefore added a second requirement that fluorescence also be elevated for the presynaptic bouton marker synapsin (see antibody table) adjacent to the PSD-95 punctum, to distinguish synapses from non-synaptic PSD-95 puncta. To further guard against misinterpretation of any non-synaptic fluorescence, we required that putative post-synaptic PSD-95 puncta persist in at least two adjacent sections, since synaptic puncta will almost always extend beyond the bounds of a single 70 nm thin section. The output of this detection process is a probability image where the value of each voxel reflects the probability that it belongs to a glutamatergic synapse. Using manually annotated synapses in the SEM channel as a baseline, we set the probability threshold at 0.75 to minimize false positives. We then merged adjacent suprathreshold voxels to define a continuous detection volume for each synapse and assigned that synapse an integer index.

### Analysis of synaptic measurements

To measure the IF signal associated with manually-annotated synapses in the conjugate IF-SEM dataset, the synapse annotations were expanded by 160 pixels (54 nm) on each side, creating a 3-D mask that typically spanned several serial sections for each synapse. The summed and the averaged signal intensity of each IF channel were computed using this mask. Synapse size was estimated in the conjugate IF-SEM dataset using the synapse contact area, calculated as the length of the postsynaptic density on each section through the synapse in the SEM channel, multiplied by the section thickness (70 nm).

For the IF-only dataset, sums and mean values of each IF channel for all voxels within the synapse detections were tabulated with one row per detection index. Synapse size in the IF-only dataset was estimated as the sum of PSD-95 immunofluorescence within the synapse detection volume as determined with the synapse detector described above. This synapse size estimate was validated using the conjugate IF-SEM dataset, which showed a strong linear relationship between synapse contact area (the ultrastructural postsynaptic density in the SEM channel and the summed PSD-95 immunofluorescence within the expanded annotation mask, Fig. 2C).

For both datasets, fluorescence values were pre-processed by first trimming outliers, excluded by criteria of lower bound < Q5 – (1.5 * (Q95 – Q5)) and upper bound > Q95 + (1.5 * (Q95 – Q5)), where Q5 is the 5^th^ percentile and Q95 is the 95^th^ percentile value for each channel. Clustering was performed after applying a log transformation to each fluorescence channel and a square-root transformation to the size; outliers were trimmed after transformation. Each column was then z-scored (Python SciKit `StandardScaler`). Clustering was performed using the Ward method and Euclidean metric (Python Seaborn `ClusterMap` function).

Silhouette analysis was based on K-means clustering with a range of 2-20 clusters (Python sk- learn `KMeans`) and subsequent silhouette analysis (Python `sk-learn.metrics silhouette_score`). Dendrograms were generated using linkage analysis (Python `Seaborn ClusterMap.Dendrogram` function). UMAPs were calculated using default settings (conjugate dataset) or by enforcing a set number of neighbors (200) and minimum distance (0.25) (Python sk-learn `UMAP` function). UMAP plots were based on 8 values: synapse size, mean fluorescence in the PSD-95, GluA1, GluA2, GluA3, GluA4, GluN1, and GluN2B channels.

In the conjugate IF-SEM dataset, the postsynaptic targets of synapses were classified as soma, dendritic shaft, dendritic spine or axon initial segment, based on their ultrastructure.

Three morphologically-defined populations of spines were analyzed: 1) “classical” spines: spines with a well-defined spine head and spine apparatus, that have characteristics typical of a strong synapse, including a perforated postsynaptic density and presynaptic mitochondria; 2) thick-necked spines: large spines with a head diameter ≥ 250 nm and neck diameter ≥ 180 nm; and 3) small spines with a head diameter < 250 nm. GABA-positive dendrites were identified by the presence of GABA-immunofluorescence and lack of spines.

In the IF-only dataset, to identify a population of synapses preferentially targeting GABA dendrites, we used the fluorescence in the GABA channel in the area surrounding the synapse. To calculate this, we dilated the synapse detection area by 6 pixels in each direction. We then selected the top 2% of synapses by GABA content.

To calculate similarity scores across synapse subclasses (Fig. 7), data were first restricted to the same set of variables used to generate UMAP plots. We then subdivided synapses based on groupings including: all synapses; putative GABA-targeting synapses (defined above); putative glutamate-targeting synapses (non-GABA); GABA- and -glutamate-targeting mix of synapses; shared postsynaptic target; shared presynaptic target; shared postsynaptic target restricted to targeting putative glutamatergic neurons; shared postsynaptic target restricted to targeting putative GABAergic neurons; synapses formed onto large (classical) spines; synapses formed onto thick-necked spines; and synapses formed onto small spines. We calculated the divergence in size and receptor content by comparing each synapse to every other synapse within the same group using linear Euclidean distance (Python `numpy` package `linalg.norm` function) on the z-scored data.

## Code availability

All code associated with this study is available online at github.com/aksimhal/single-synapse-proteomics.

## Abbreviations

AMPAR: The α-amino-3-hydroxy-5-methyl-4-isoxazolepropionic acid receptor
NMDAR: The *N*-methyl D-aspartate receptor
GABA: gamma-aminobutyric acid
EM: Electron microscopy
SEM: Scanning electron microscopy
IF: Immunofluorescence
AT: Array tomography
PSD-95: Postsynaptic density protein 95
TeA: Temporal association cortex
UMAP: Uniform manifold approximation and projection

## Extended data

**Extended Data Figure 1.**
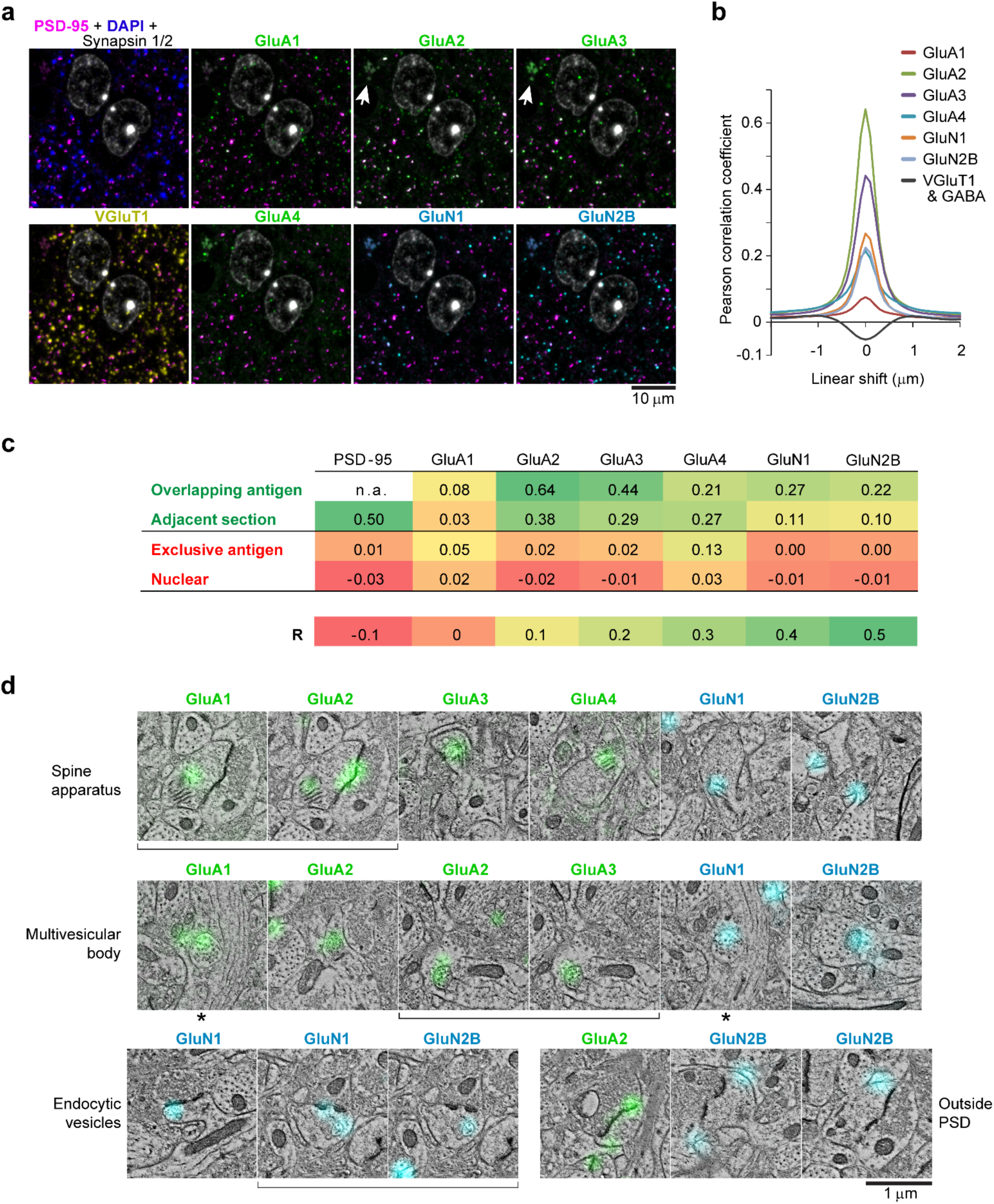
Immunofluorescence labeling of AMPA and NMDA receptor subunits. **a.** Single 70 nm section showing colocalization of presynaptic (synapsin and VGluT1) and postsynaptic immunolabels (AMPA and NMDA receptor subunits) with PSD-95 (magenta). Nuclei are labeled with DAPI (white). Arrow points to lipofuscin granules which are autofluorescent with broad emission spectrum and visible in all channels (arrow only shown in two of the channels). **b.** Pearson correlation coefficient as a function of the lateral offset between each glutamate subunit receptor channel and PSD-95. This controls for random overlap between two antibody labels^1^. When there is true correlation, the Pearson correlation coefficient is highest with no shift between the channels, and quickly drops to zero (no correlation) as the channels are shifted laterally relative to each other. All glutamate receptor subunit antibodies show correlation with PSD-95. When two channels are anticorrelated, as the case of VGluT1 and GABA, there is negative correlation at no shift and it gradually increases with lateral offset. **c.** The performance of the glutamate receptor subunit antibodies is evaluated using the Pearson correlation coefficients from 4 different control comparisons. As a reference, the PSD-95 antibody is an excellent antibody that specifically labels the postsynaptic density of glutamatergic synapses in neocortex. Colocalization (green) is expected in the top two tests, and random distribution or anticorrelation (yellow/red) are expected in the 2 tests below. These tests reveal that GluA1, even though it colocalizes with PSD-95, is the least sensitive antibody. The GluA2, 3 and 4 antibodies perform very well, and the lower R value for GluA4 in the overlapping antigen test is expected due to its much more restricted distribution at adult neocortical synapses. The GluN1 and N2B antibodies also perform as expected, given that they are enriched at smaller synapses that span only one or two sections, which explains their lower Pearson coefficient in the adjacent sections test. *Overlapping antigen test* for the specificity of staining: The overlap between glutamate receptor subunits and PSD-95 is reported here, identical to the Pearson coefficient with no shift shown in b. Less abundant targets give lower coefficients because they are present at a smaller subset of postsynaptic densities. *Adjacent section* test for the consistency of immunolabeling: Because the distribution of targets is very similar on two adjacent ultrathin sections (70 nm thickness), colocalization of immunolabeling between adjacent sections is expected. This correlation is influenced by antibody characteristics, but also the size of targets, with smaller targets displaying larger spatial variability from section to section. *Exclusive antigen* test for antibody specificity: The overlap of the tested antibodies with the inhibitory neurotransmitter GABA is reported here. Values of R around 0 are expected. The high R value for GluA4 and to a lesser extent GluA1, is likely due to the preferential distribution of these subunits at synapses onto GABA targets, thus resulting in adjacency of GluA4 / GluA1 and GABA. *Nuclear* test: All antibodies were also compared with DAPI to control for background nuclear immunolabeling. Values of R around 0 are also expected here. **d.** Receptor immunolabeling in the spine apparatus, multivesicular bodies within dendrites, endocytic vesicles in spines, and plasma membrane adjacent to the postsynaptic density. Brackets indicate pairs of images that are from the same section of the same synapse, and asterisks indicate adjacent sections from the same synapse.

**Extended Data Figure 2.**
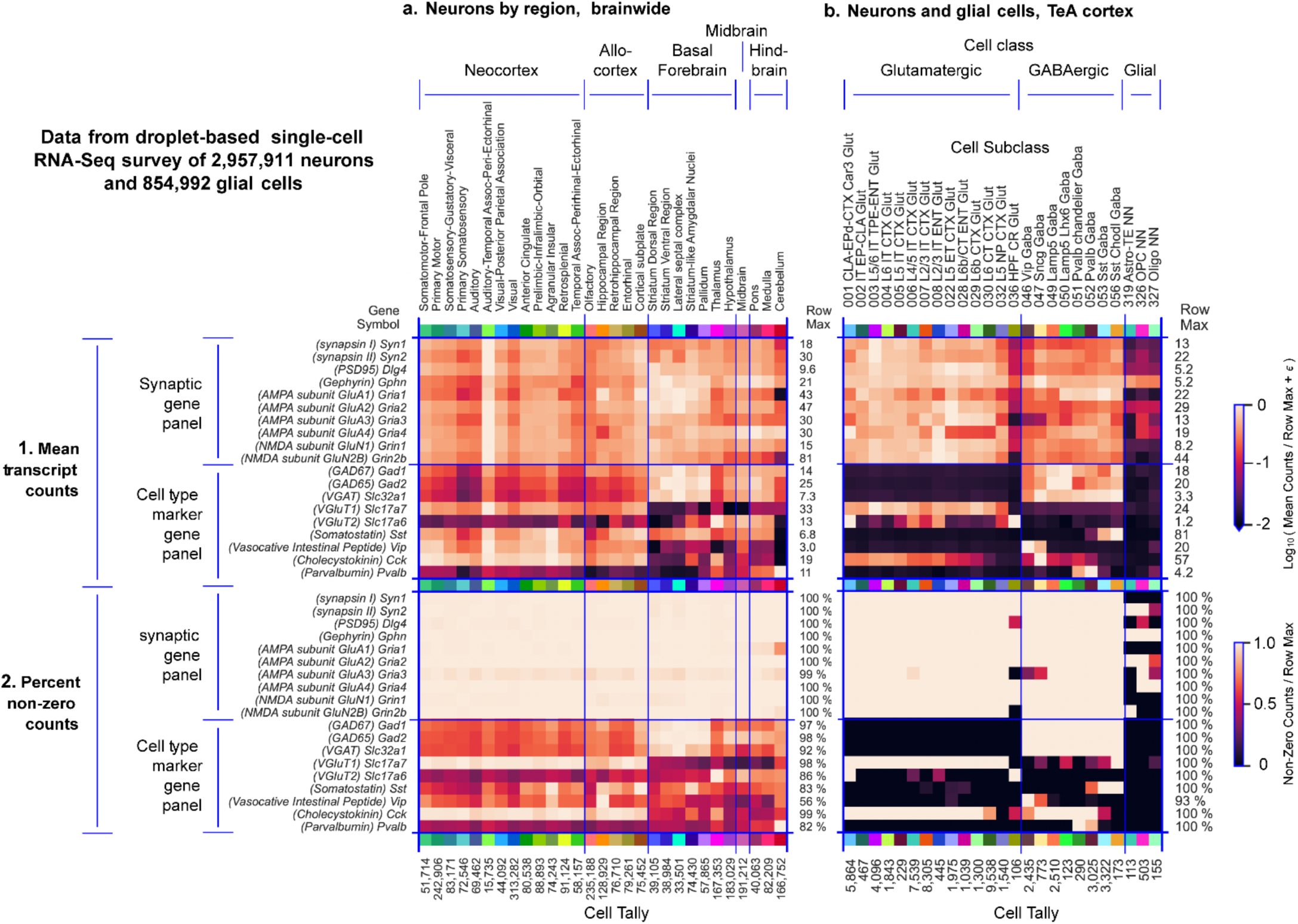
Single-cell mRNA-Seq data show the expression of proteins targeted by our synaptic antibodies throughout the mouse brain, including temporal association (TeA) cortex. **a, b.** Transcriptomic analysis using the Allen Institute’s single-cell RNA-Seq data showing the transcriptomic counts of the immunomarkers used in this study across cell types in temporal association cortex. Data resources and cell type taxonomies are described elsewhere^2^. Heat maps represent mean mapped counts (row group 1) and fraction of cells with non-zero mapped counts (row group 2) by gene per brain region, brain wide (column group **a**) or per cell subtype in TeA region (column group **b**). Maps in both a and b quantify mRNA-level expression of 10 genes encoding 10 proteins targeted by our synaptic antibody panels and 8 commonly recognized cell-type markers. The marker gene panels are included to contrast the highly differential expression of the cell-type markers with the much more uniform expression of synaptic panel genes. Results from glial cells are included in b to contrast the subtype-restricted expression of the synaptic panel genes in glia with the very broad expression of same in neurons. Python analysis scripts used to generate heat maps in this figure are publicly available (github.com/aksimhal/single-synapse-proteomics).

**Extended Data Figure 3.**
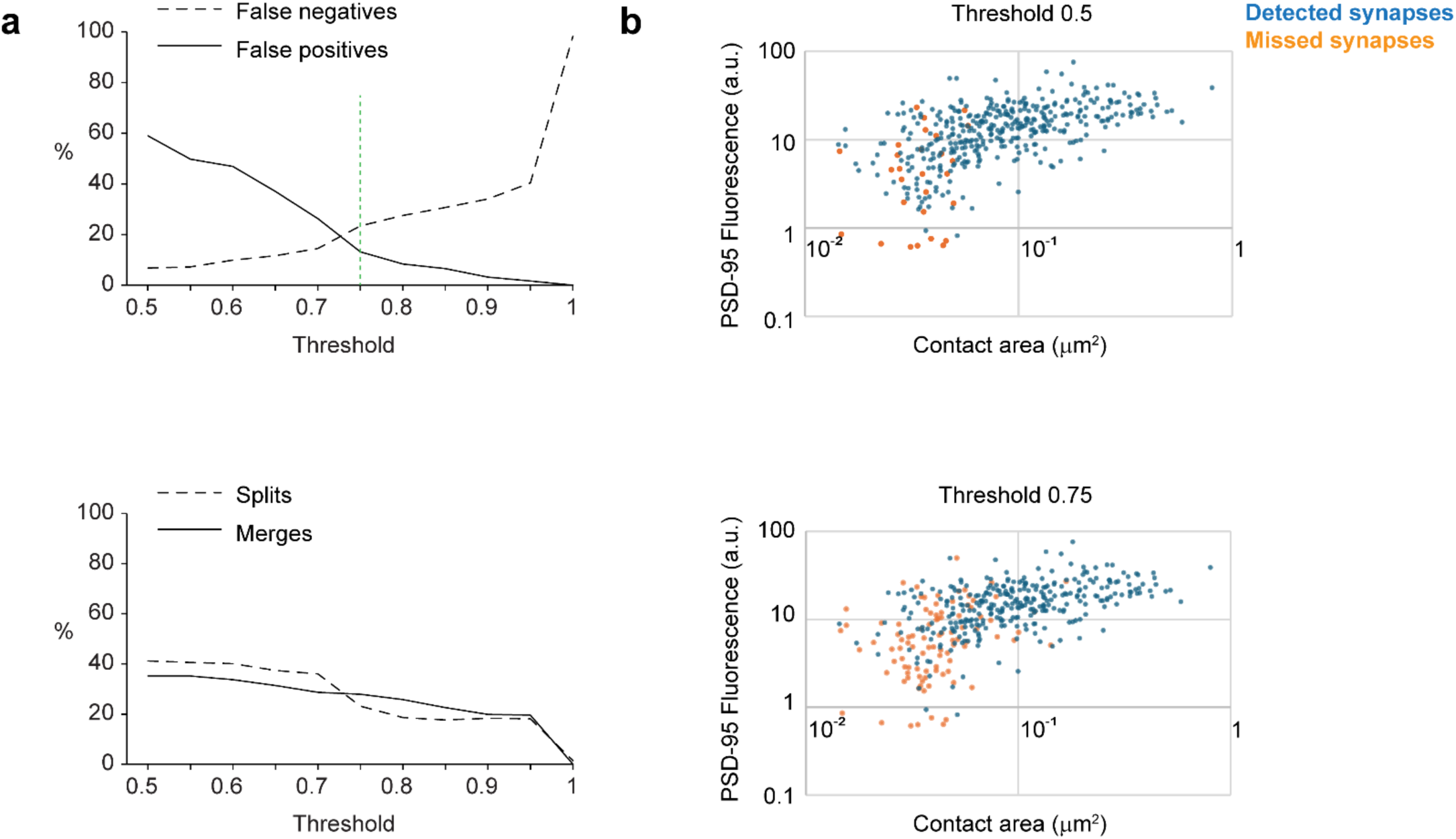
Ultrastructural evidence supports IF-based synapse detection and analysis. **a.** Further evaluation of our probabilistic synapse detection approach by comparing the automated synapse detection and manual annotation of synapses in the conjugate dataset. The query for synapse detection was defined as a PSD-95 punctum on two consecutive sections and an adjacent synapsin punctum present on at least one section. Plots of false positives and false negatives (top), and splits and merges (bottom) as a function of threshold. Based on these measurements, a threshold of 0.75 was chosen for synapse detection in the IF dataset (green dashed line). **b.** Scatterplot illustrating the synapse contact area as estimated on the electron micrograph and PSD-95 fluorescence of detected and missed synapses using 2 different thresholds for automated synapse detection. Undetected synapses are generally small and with low PSD-95 fluorescence.

**Extended Data Figure 4.**
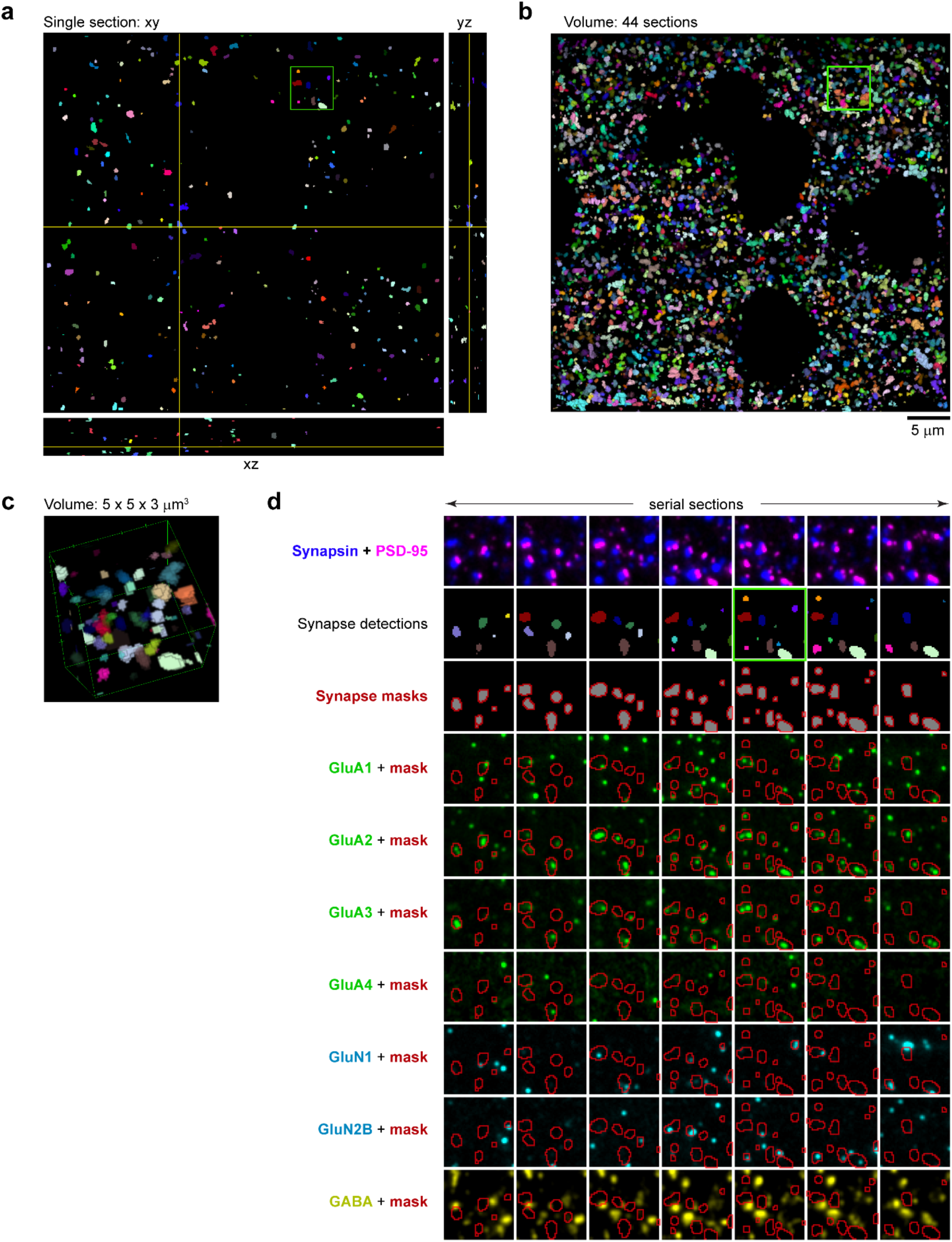
IF-AT enables the measurement of multiplex immunofluorescence at thousands of individual synapses. **a.** Synapses detected within the IF-only volume using the probabilistic synapse detector with a query of PSD-95 on 2 consecutive sections and adjacent synapsin present on 1 section, and threshold of 0.75 probability. Each synapse is displayed with a different color. The volume is sectioned in the xy, xz and zy plane. **b.** The same synapse detections presented as a volume. **c.** The area within the small green square in a and b, presented as a volume. **d.** 7 consecutive sections from the volume in c, showing the immunofluorescent channels used for synapse detection in the top row, followed by the synapse detections (2^nd^ row), then the masks where the fluorescence intensities were calculated for each synapse (3^rd^ row). The area of the mask is equal to the area of the detection. The remaining rows show the overlap of the synapse masks with the immunofluorescence for each receptor subunit, and with GABA.

**Extended Data Figure 5.**
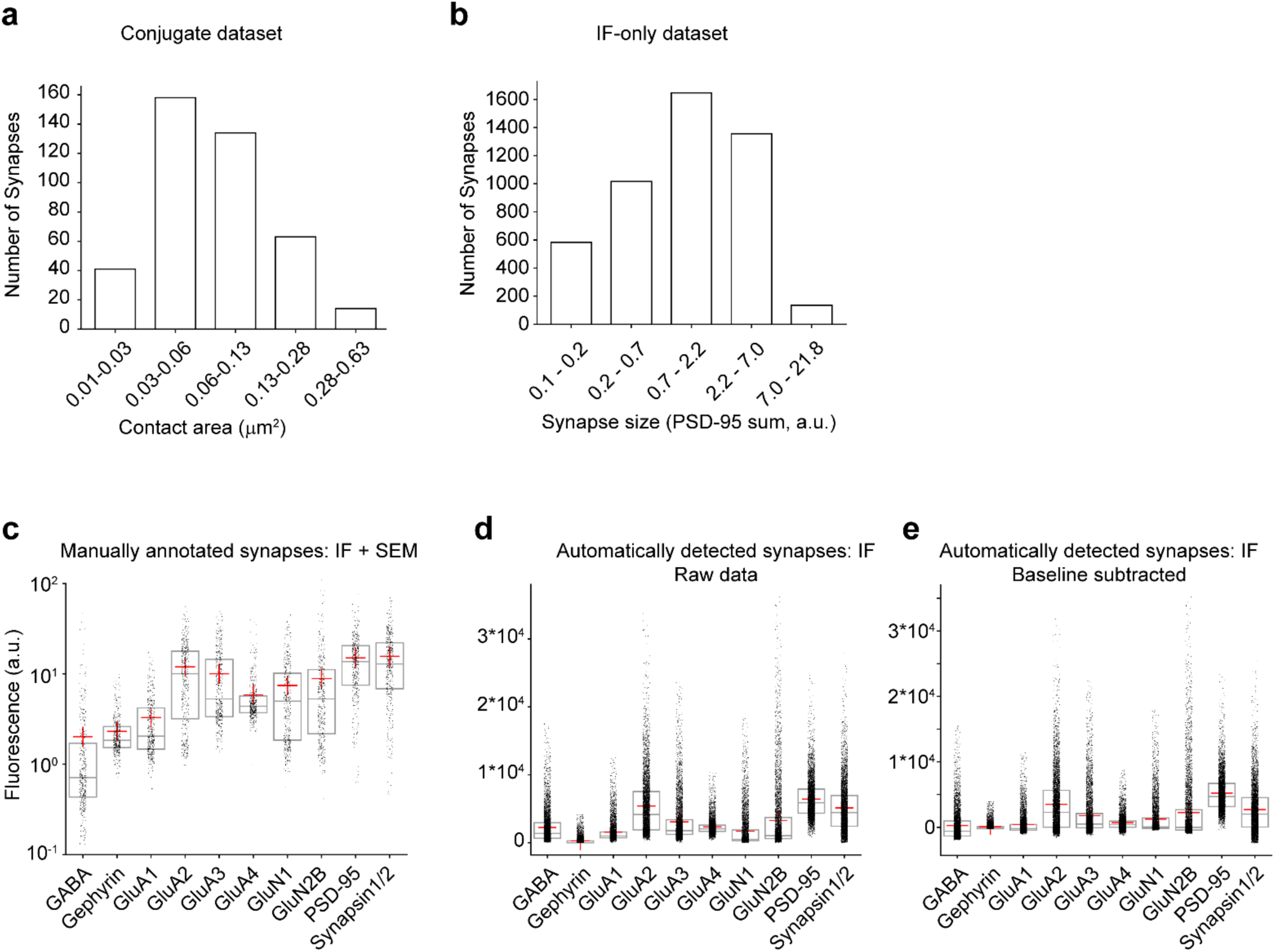
IF-based synapse analysis yields results comparable to conjugate AT. **a.** Size distribution of synapses from the conjugate dataset. Synapses are binned by contact area, estimated by the presence of a postsynaptic density in the scanning electron micrographs. **b.** Size distribution of synapses from the IF-dataset. Synapses are binned by summed PSD-95 fluorescence used as an estimate for synapse size. **c.** Average immunofluorescence at manually annotated synapses within the conjugate dataset. **d.** Average immunofluorescence at automatically detected synapses within the IF dataset. **e.** Same as d, with the baseline immunofluorescence subtracted. To define baseline fluorescence, we shuffled the synapse detections by 180° rotation in the XY plane and 50% translation along the Z-axis.

**Extended Data Figure 6.**
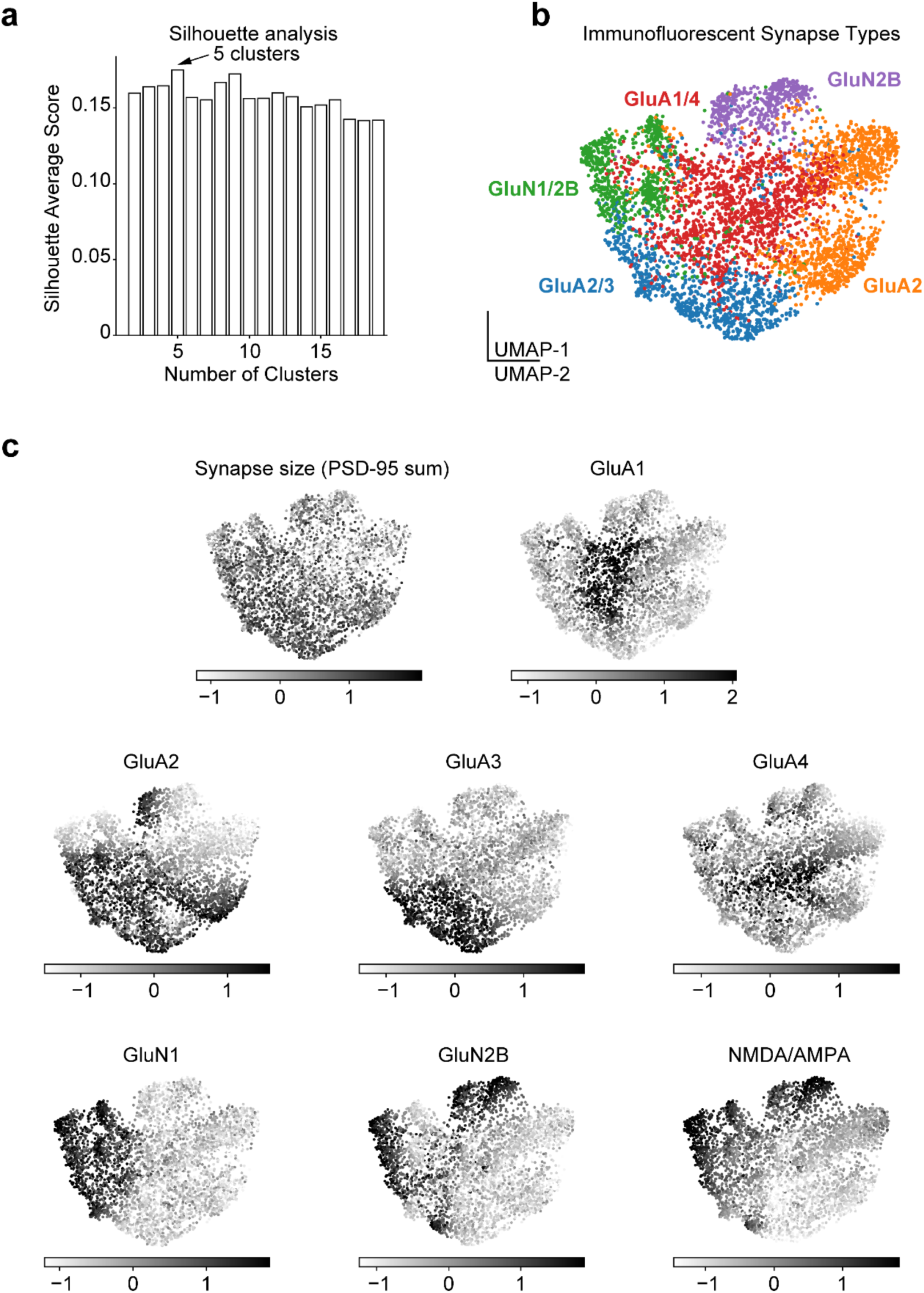
Immunofluorescent synapse subclasses. **a.** Silhouette analysis identifies 5 distinct clusters in the IF-only dataset. **b.** UMAP dimensionality reduction plot, colored by clusters defined in the main Fig.2e reveals clustering of distinct glutamatergic synapse subclasses in the IF-only dataset. Also included in main Fig.2f. **c.** Grayscale coding of UMAP projections by synapse size (summed PSD-95 immunofluorescence), glutamatergic receptor expression (average immunofluorescence of GluA1 - 4, GluN1, GluN2B), and NMDA/AMPA ratio in the IF-only dataset. Four of the c panels are also included in the main Fig.2j.

**Extended Data Figure 7.**
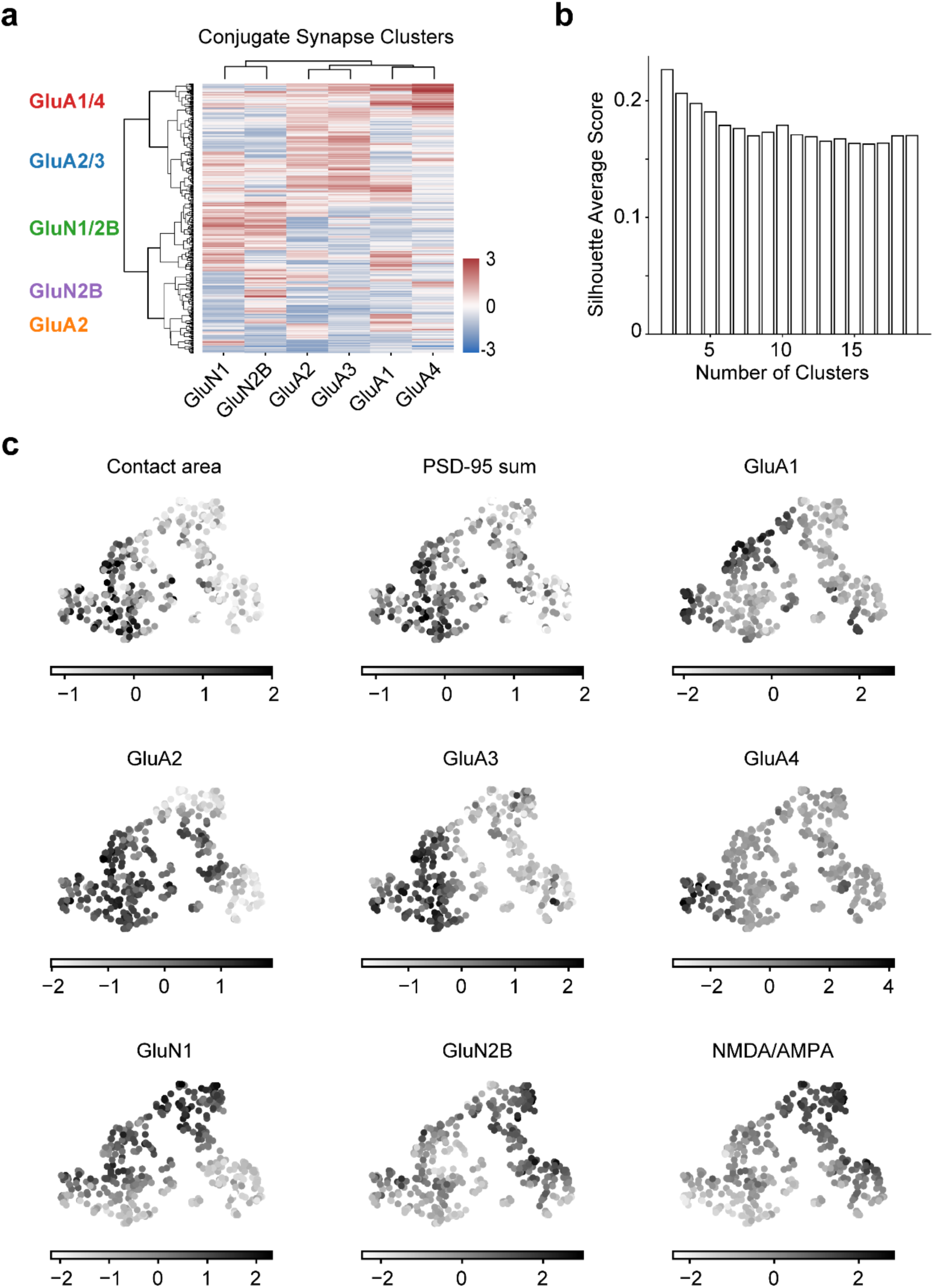
Conjugate synapse subclasses. **a.** Automated clustering of glutamatergic synapses from the conjugate dataset based on synapse immunolabeling characteristics (n=410 synapses). Similar synapse groupings as in the IF-only dataset are indicated to the left (main Fig.2e). **b.** Silhouette analysis does not show a clear peak in the conjugate IF-SEM dataset, likely due to the low number of synapses. **c.** Grayscale coding of UMAP projections by synapse contact area, summed PSD-95 immunofluorescence and average glutamatergic receptor immunofluorescence in the conjugate IF-SEM dataset. Four of the c panels are also included in the main Fig.7g.

**Extended Data Figure 8.**
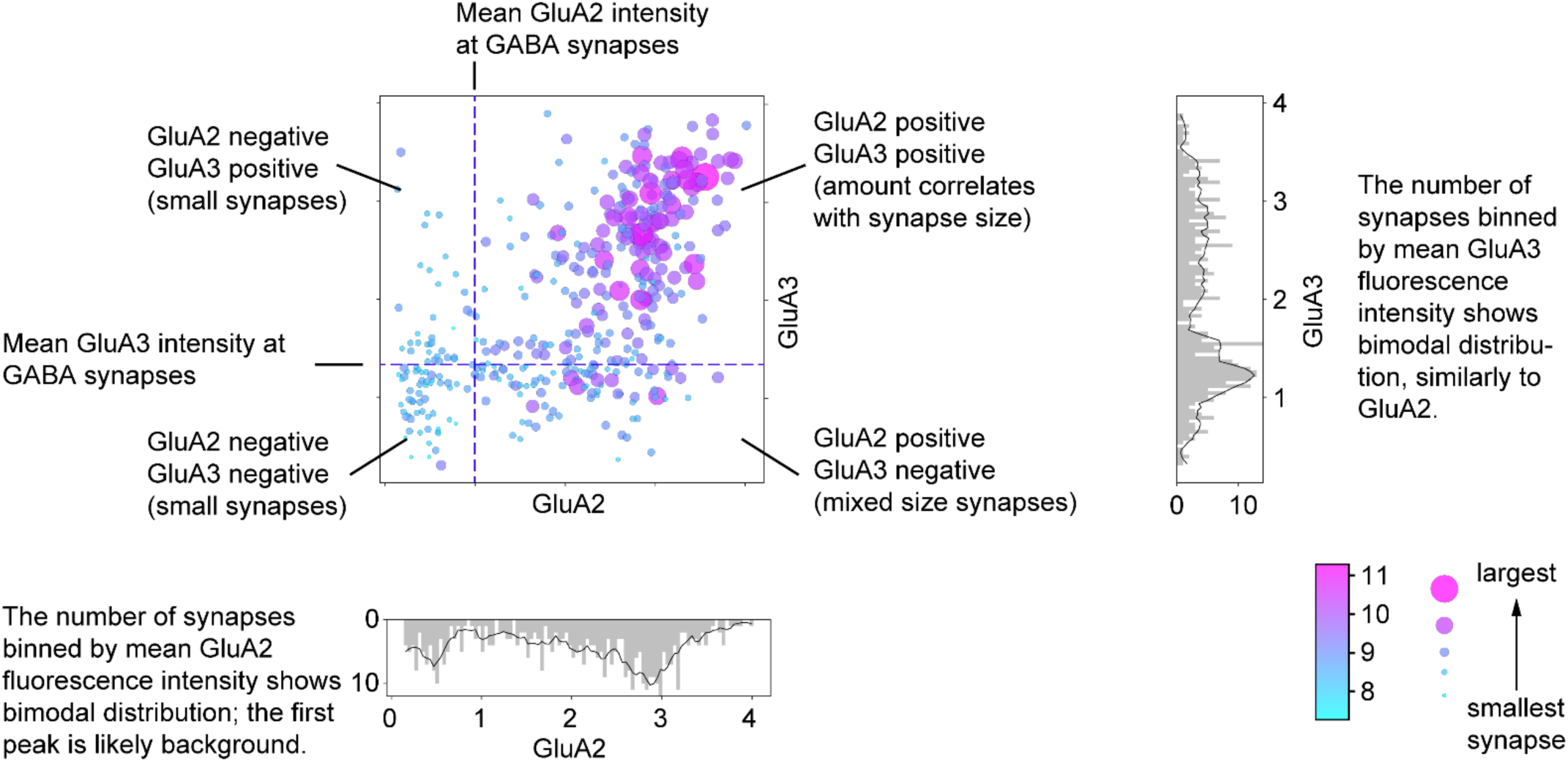
Guide to scatterplots. Scatterplot comparing mean intensity of the GluA2 and GluA3 fluorescence channels at annotated synapses (conjugate IF-SEM dataset). Same as in main Fig.3f. This scatterplot is used to provide a guide to reading and interpreting the scatterplots in the main Fig.3 and Fig.5.

